# geneRNIB: a living benchmark for gene regulatory network inference

**DOI:** 10.1101/2025.02.25.640181

**Authors:** Jalil Nourisa, Antoine Passemiers, Jeremie Kalfon, Marco Stock, Berit Zeller-Plumhoff, Robrecht Cannoodt, Christian Arnold, Alexander Tong, Jason Hartford, Mihai G. Netea, Antonio Scialdone, Laura Cantini, Yves Moreau, Daniele Raimondi, Yang Li, Malte D. Luecken

## Abstract

Gene regulatory networks (GRNs) underpin cellular identity and function, playing a key role in health and disease. GRN inference has received substantial attention, motivating systematic benchmarking. Despite various benchmarking efforts, existing studies remain limited in the number of methods, datasets, and metrics, fail to capture the context-specific nature of regulatory interactions across biological conditions, and are constrained by the absence of a reliable ground truth. Here, we introduce geneRNIB, a comprehensive GRN inference benchmarking framework built on three key principles: continuous integration, context-specific evaluation, and holistic assessment in the absence of a true reference network. geneRNIB enables the seamless incorporation of new algorithms, datasets, and evaluation metrics to reflect ongoing developments. In the current version, we systematically integrated and assessed 12 GRN inference methods, spanning single- and multiomics approaches across 11 datasets including thousands of perturbation scenarios. We introduced complementary metrics specifically designed to assess context-specific inference. Our findings indicate that simple models with fewer assumptions often outperform more complex pipelines across several perturbation-informed and predictive metrics. Notably, gene expression-based algorithms yielded better results than more advanced multimodal approaches. In addition, we identify several potential factors that influence the performance of GRN inference and offer actionable guidelines for the future development of the method. By addressing these critical limitations in existing benchmarks, geneRNIB advances GRN inference research and fosters progress toward personalized medicine.

## Introduction

Gene regulatory networks (GRNs) are fundamental to understanding cellular identity and behavior in health and disease. GRNs describe how gene expression is controlled through complex interactions between transcription factors (TFs), cis-regulatory elements (CREs), cofactors, as well as epigenetic and trans-regulatory mechanisms [1]. Given the cost and time constraints of direct experimental validation of regulatory interactions, computational GRN inference has become an increasingly valuable approach. However, GRNs exhibit context-specific variations depending on the organism, cell type, disease state, and environmental conditions [2, 3], requiring GRN inference methods to model regulatory dynamics for the given condition. Moreover, different computational approaches often yield inconsistent results, and their accuracy remains difficult to validate due to the limited availability of experimentally confirmed interactions [4, 5, 2]. To address this, continuous benchmarking is necessary, not only to integrate new GRN inference methods, but also to refine evaluation metrics and datasets, ensuring robust and reliable inference across diverse biological contexts.

The advent of single-cell sequencing technologies has transformed GRN inference by drastically increasing the size of data, offering single-cell resolution, and allowing reconstruction of context-specific regulatory networks [3, 6, 7]. A persistent challenge in GRN inference is causal inference, driving the exploration of diverse computational and experimental approaches for improvement [8, 2, 9]. The integration of interventional data, such as gene knockouts, longitudinal data, and multimodal datasets, has been explored to enhance causal discovery [8, 10, 6, 2]. Given that regulatory interactions span multiple molecular layers—including DNA, RNA, and chromatin modifications—there has been increasing interest in multiomics approaches for GRN inference [11, 3, 12, 5]. In particular, integration of gene expression and chromatin accessibility data has shown promise due to its cost efficiency and its potential to improve causal inference [2, 3, 12].

Benchmarking GRN inference is complicated by the lack of ground truth data, as direct measurement of regulatory interactions at high throughput remains infeasible. To address this, various strategies have been proposed. One approach involves simulated networks with synthetic gene expression data [13, 14]. Although these data sets provide controlled scenarios for evaluation, they are often not representative of true biological signals [15, 16]. Another approach relies on proxy measures of ground truth, known as silver standards [2, 13], by defining a reference with experimental priors derived from diverse sources, such as TF binding data, perturbation experiments, or curated databases from the literature and public repositories [13, 17, 18].

Although valuable, these resources are inherently noisy and capture only a fraction of known interactions. In addition, their diverse origins further obscure context-specific regulatory dynamics, making it challenging to derive precise, condition-specific insights. Moreover, TF binding alone does not guarantee regulatory activity, limiting their effectiveness as a ground truth for gene regulation. Alternatively, indirect approaches have been designed to bypass the reliance on ground truth. For instance, the CausalBench challenge introduced multiple metrics to evaluate the causality of inferred TF-target gene interactions by comparing changes in target gene expression following perturbations to TFs [8]. Kamal et al. [5] proposed a feature-based evaluation framework that builds regression models on top of the feature space defined by the GRN topology. The core assumption of this approach is that GRNs capturing biologically meaningful regulatory structure induce more informative feature representations, which in turn yield better predictive performance. Badia et al. recently introduced `GRETA`, a modular framework (i.e., enabling the systematic evaluation of individual analytical steps) that integrates multiple evaluation metrics based on both prior knowledge and indirect approaches [4].

While these studies provide valuable and complementary benchmarking insights, they are inherently shaped by data availability and their specific design choices. For example, the CausalBench challenge focuses on a limited number of datasets available at the time and on selected few proxy evaluation metrics, enabling controlled comparisons across methods. In contrast, Badia et al. benchmark multimodal GRN inference using a broad range of metrics and datasets. To support this scope, they aggregate data across multiple contexts, which resulted in perturbation data being derived from experiments distinct from those used for network inference, thereby reducing context specificity. As a result, these studies emphasize different aspects of GRN benchmarking and highlight the trade-offs between controlled large-scale evaluation and the ability to capture biological constraints. By design, any benchmarking effort is constrained in its inclusion of methods, datasets, and evaluation strategies, particularly as technologies and data resources continue to evolve. Therefore, an adaptable benchmarking framework is essential for continuously integrating new developments and establishing robust criteria for the selection and advancement of GRN inference methods.

Here, we present geneRNIB (gene Regulatory Network Inference Benchmark), an open-source and fully reproducible platform for benchmarking GRN inference. geneRNIB introduces three major advances; (i) It is a community-driven, continuously evolving benchmark designed to incorporate new datasets, metrics, and inference algorithms as they become available; (ii) geneRNIB emphasizes context-specific evaluation of GRN inference by controlling for cell type and experimental variability; (iii) geneRNIB offers standardized evaluation metrics that account for variability across datasets and GRN models, promoting robust and fair comparisons across methods. It leverages the largest single-cell perturbation compendium to date, comprising more than 13 million cells and a broad range of perturbation types, to ensure that GRN inference and assessment are performed within the same biological context. Finally, the benchmark supports both transcriptomics-based and multiomics GRN inference methods. Using geneRNIB, we show that GRN inference performance depends strongly on biological context, with intrinsic dataset properties—such as gene expression level, variability, and perturbation strength—substantially shaping outcomes. Under context-matched, standardized evaluation, simple expression-based methods match or outperform more complex multimodal approaches. We show that identified top-performing models yield regulatory interactions that are systematically more accurate across genes and TFs and are more stable across perturbations and donors. In addition, these methods recover regulatory circuits in a multiomics dataset of multiple myeloma precursor (MMP) disease that are missed by low-performing models, demonstrating that geneRNIB recommendations directly translate to real-world settings. We envision that geneRNIB will pave the way toward standardized, context-aware, and statistically rigorous evaluation of GRN inference methods, providing guidance to data analysts and method developers while improving our understanding of regulatory interactions.

## Results

### A living benchmarking framework for context-specific GRN inference

Gene regulatory interactions occur in specific biological contexts, shaped by factors such as cell type, disease, and experimental condition. To evaluate GRN inference methods within the contexts in which regulatory interactions are inferred, we developed geneRNIB, a living benchmarking framework for context-specific evaluation (Fig. 1). Unlike static benchmarks, geneRNIB automatically re-evaluates previously assessed methods and updates the leaderboard as new components are introduced, ensuring reproducibility and upto-date performance tracking. In its current form, geneRNIB integrates 11 datasets, 12 GRN inference methods, and 11 evaluation metrics. It is hosted on *Open Problems in Single-Cell Analysis* ^1^, a communitydriven platform that leverages modern containerization and workflow technologies—including `Docker`, `Viash`, and `Nextflow`—together with cloud infrastructure to ensure scalability and accessibility. This design enables the integration of new methods, datasets, and metrics, maintaining adaptability to evolving community standards and ensuring the long-term relevance of the framework (Supplementary Note 1).

**Figure 1:**
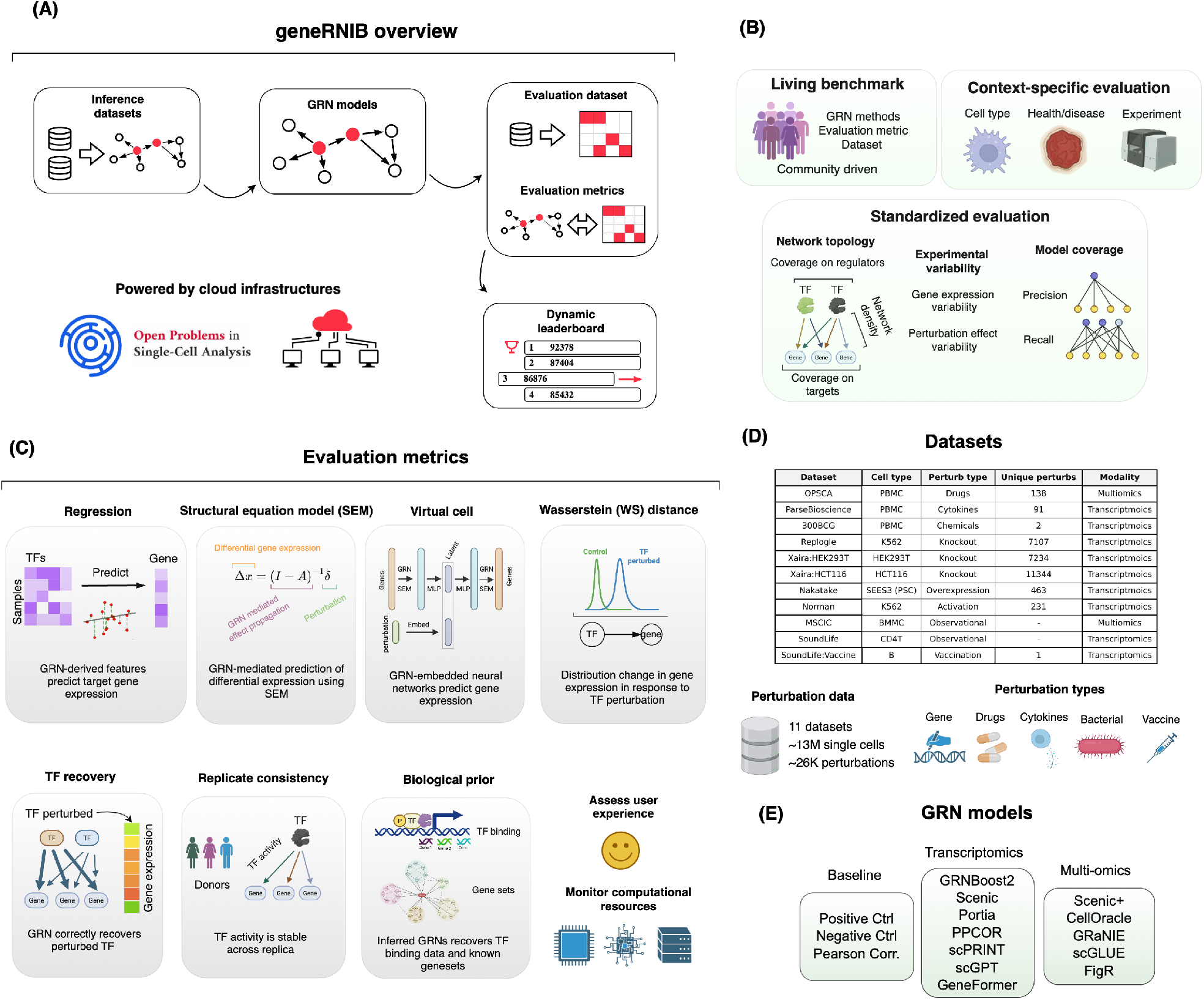
Overview of geneRNIB. **(A)** geneRNIB provides a reproducible, cloud-hosted computational infrastructure that integrates inference and evaluation datasets, GRN models, standardized evaluation metrics, and a dynamic leaderboard to track state-of-the-art GRN benchmarks. **(B)** geneRNIB is a communitydriven, continuously evolving benchmark that supports future integration of new GRN inference methods, evaluation metrics, and datasets. It enables context-specific assessments across cell types, conditions, and experimental settings, and ensures fair comparison through standardized metrics that account for confounding factors such as network topology differences, experimental variability, and GRN model coverage by reporting both precision and recall. **(C)** Overview of evaluation metrics used for GRN evaluation. The three predictive metrics—Regression, SEM, and Virtual cell—evaluate how well the TF-gene relationships given in GRN models predict diverse perturbational gene signatures. Other metrics provide orthogonal evaluation of GRN inference. **(D)** Overview of datasets used in geneRNIB. The resource compiles 11 large-scale datasets totaling around 13 million single cells and roughly 25,000 unique perturbations. These datasets span diverse perturbation types, including bacterial stimuli, chemical and drug treatments, cytokine exposures, gene knockouts, and gene overexpression. **(E)** GRN models integrated in geneRNIB. Multiomics GRN inference methods integrate scRNA-seq and scATAC-seq, leveraging an enhancer-driven approach to identify regulatory interactions. In contrast, transcriptomics-based methods rely solely on gene expression data to infer regulatory networks. Additionally, three baseline models—Positive Control, Negative Control, and Pearson Correlation—were implemented as reference benchmarks. Some elements of the figure were created with BioRender.com.

Given the absence of definitive ground truth GRNs, we developed multiple metrics to evaluate GRN inference, organized into two main categories: predictive metrics and orthogonal validation metrics (Fig. 1-C and Methods). The predictive metrics assess whether inferred GRNs can accurately predict cellular responses to perturbations. The Regression metric evaluates whether the regulators identified by a GRN can accurately predict the expression of target genes. The Structural Equation Model (SEM) metric is based on a causal model of steady-state gene expression, where each gene’s expression is modeled as the result of regulatory interactions with other genes, as defined by a GRN model. The Virtual cell metric integrates GRN structures into neural network architectures to predict unseen cell type/perturbation pairs. To complement these approaches, we incorporated additional metrics that provide orthogonal validation based on distinct biological assumptions and experimental evidence. These include the Wasserstein Distance (WS) metric, which quantifies shifts in target gene expression distributions following perturbations of upstream TFs; the TF recovery metric combines perturbation data with inferred network edges to identify the perturbed TF in each experiment; the Replicate Consistency metric evaluates the reproducibility of inferred TF–gene relationships by measuring the stability of TF activity patterns across biological or technical replicates; the TF binding metric compares inferred networks against experimentally validated TF–target interactions from ChIP-seq databases; and the Gene sets recovery metric evaluates whether the inferred GRN captures biologically meaningful functional relationships by assessing the enrichment of canonical pathway gene sets among target genes. In total, we implemented 8 main metric groups, expanding to 11 individual metrics when considering both precision and recall for Regression, WS Distance, and TF recovery. Among these metrics, SEM, Virtual cell, and Replicate Consistency were newly developed for this benchmark, while the remaining metrics were adopted from previous studies (Methods).

To ensure fair and comparable evaluation, we implemented several key improvements across metrics. These include (i) edge selection strategies to account for differences in network density across methods, (ii) baseline controls to account for perturbation-specific biases and establish expected performance ranges, and (iii) precision- and recall-like formulations to capture both high-confidence predictions and network coverage. Together, these refinements enable robust comparison of GRN inference methods under diverse conditions.

To evaluate methods across a wide range of biological contexts, we curated 11 benchmark datasets spanning diverse perturbations and cellular systems (Fig. 1-D and Methods). Nine datasets—Replogle, Nakatake, Norman, Xaira-HEK293T, Xaira-HCT119, 300BCG, SoundLife, SoundLife:Vaccine, and ParseBioscience—support transcriptomic GRN inference, while OPSCA and MSCIC enable multiomics GRN inference by integrating paired transcriptomic and chromatin accessibility data. These datasets encompass observational and longitudinal data, as well as diverse perturbation modalities (bacterial stimulation, gene knockouts/knockdowns, chemical drugs, cytokines, and vaccination), totaling approximately 13 million cells and 26,000 unique perturbations (Extended Data Table 1).

Finally, geneRNIB incorporates 12 GRN inference methods (Fig. 1-E and Methods). These include three baseline methods (Positive Control, Negative Control, Pearson Correlation), seven transcriptomics-based methods (GRNBoost2 [19], Scenic [10], Portia [9], PPCOR [20], scPRINT [21], Geneformer [22], scGPT [23]), and five multiomics methods (Scenic+ [12], CellOracle [3], GRaNIE [5], scGLUE [11], FigR [24]). The Pearson Correlation baseline provides a simple co-expression reference using transcriptomic data alone. The Positive Control, which leverages both inference and evaluation data, establishes an approximate upper bound on achievable performance, while the Negative Control, based on randomized networks, defines a lower bound.

Overall, geneRNIB is a large-scale and diverse benchmark that captures a broad spectrum of evaluation criteria for GRN inference and is designed to remain extensible through continued community contributions.

### Evaluation metrics are context specific and robust to spurious correlations and preprocessing choices

We systematically evaluated whether our metrics reliably capture valid regulatory relationships through a series of experiments designed to test their robustness to context specificity, structural properties of GRNs, causality, and confounding data preprocessing or modeling choices. To test whether our evaluation metrics capture context-dependent regulatory patterns, we compared GRN models inferred from our datasets with models derived from heterogeneous cell types and tissues from external sources (Methods). Because the evaluation data are intrinsically linked to the data used for GRN inference, we expected our inferred models to achieve higher performance. Indeed, across multiple Peripheral Blood Mononuclear Cells (PBMC)-derived datasets, including OPSCA, ParseBioscience, 300BCG, and SoundLife datasets the inferred GRN models consistently outperformed those sourced from elsewhere (Fig. 2-A and Supplementary Fig. 3). These findings indicate that the inferred GRNs capture regulatory relationships specific to the cell type, experimental conditions, and biological context, and that our evaluation metrics are successful in context-specific evaluation.

**Figure 2:**
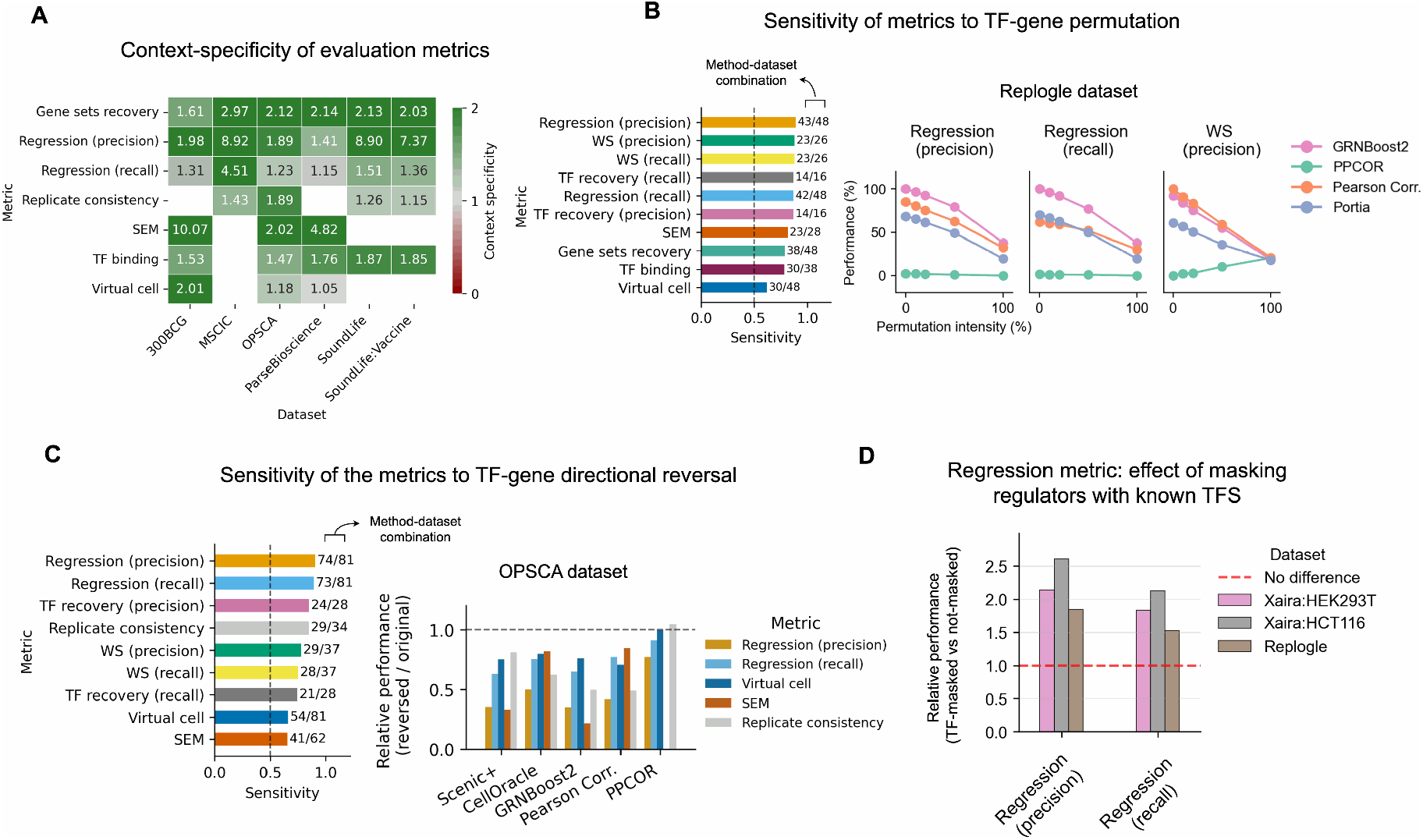
Context specificity and robustness of the evaluation metrics. **(A)** Sensitivity of the metrics to context-specific GRNs. Performance of GRN models inferred from context-matched datasets is compared with GRNs derived from unrelated experiments, cell types or tissues. Values represent the relative mean performance of inferred GRNs versus non-specific GRNs. **(B)** Permutation experiments of TF–gene connections. Sensitivity is defined as the relative mean performance of original GRNs compared with permuted GRNs, where higher values indicate a larger performance drop after permutation (left). Bars are annotated with the number of dataset-method combinations for which the metric showed a decrease in performance after permutation. The (right) panel shows example dataset–metric combinations illustrating the progressive performance decline with increasing permutation intensity. Of note, the increased trend in PPCOR indicate a below baseline performance of this model. **(C)** Permutation experiments of TF–gene directionality. Sensitivity values across dataset–method combinations are shown on the (left), while example dataset–metric–method combinations are illustrated on the (right). **(D)** Performance comparison of GRN inference using the Regression metric with and without restricting candidate regulators to known TFs.

Second, to assess whether the metrics are influenced by spurious correlations, we applied arbitrary permutations (Methods) to TF–gene connections and observed a consistent decline in performance with increasing degrees of permutation (Fig. 2-B and Supplementary Fig. 4). Notably, metrics such as Regression and WS distance showed a performance drop even at 10% permutation intensity. This indicates that inferred TF–gene regulatory interactions better generalize across unseen perturbations, suggesting that the models capture functionally relevant relationships that reflect underlying regulatory structure rather than spurious associations. Furthermore, our metrics are sensitive to the progressive degradation of these networks under increasing permutation. However, permuting regulatory weights (regulatory confidence) and interaction signs (positive versus negative) did not produce a notable decline in performance (Extended Data Fig. 1-B and Supplementary Fig. 4), indicating that the metrics are less sensitive to these properties.

To evaluate whether our metrics prioritize causal relationships over pure correlations, which remains a central challenge in GRN benchmarking, we conducted two experiments (Methods). First, a distinguishing feature of causal GRNs is the directionality of edges, a property that purely correlational models lack. To assess this, we systematically reversed edge directionality (e.g., flipping *A* → *B* relationships to *B* → *A*) and observed a substantial decline in performance across most metrics (Fig. 2-C, Extended Data Table 1-E). Notably, the Regression, TF recovery, and WS distance metrics showed the highest sensitivity to edge reversal. TF binding and Gene set recovery metrics were not included in this experiment because reversing TF–gene relationships violates the assumptions underlying these metrics (see Supplementary Note 3).Second, we further evaluated the Regression metric by comparing its performance under two strategies: (a) restricting candidate regulators to known TFs in gene-wise model construction, and (b) selecting regulators solely based on high co-expression without prior filtering. TF-filtered models consistently achieved higher scores than the baseline models (Fig. 2-D). Taken together, these two experiments indicate that our metrics, grounded in perturbation data, prioritize causal relationships (direct or indirect) over pure correlations. This is supported by theoretical evidence that causal relationships exhibit greater invariance across interventions than correlational associations [25]. For instance, perturbations may disrupt co-expression patterns while leaving the underlying causal regulatory structure intact.

To ensure the Regression metric is not confounded by model choice, we compared Ridge regression (our default due to computational efficiency) with a nonlinear gradient boosting model (GBM). The relative performance of GRN methods was highly consistent between the two approaches (Extended Data Fig. 1-C), confirming that the metric preserves comparative conclusions regardless of regression algorithm. Finally, considering that preprocessing, especially normalization, may influence GRN inference, we evaluated the consistency of benchmark scores under different normalization approaches (Methods). Specifically, we repeated GRN inference and evaluation using two normalization methods for both inference and evaluation datasets, namely shifted logarithmic normalization (default) and Pearson residuals. The comparative performance remained highly consistent across both normalization methods (Extended Data Fig. 1-D). For example, the top-performing models (GRNBoost2 and Pearson Correlation) remained top performers across both normalization methods. This suggests that the inferred regulatory relationships—rather than normalization artifacts—drive the performance in our benchmark.

Collectively, these experiments establish the proposed evaluation metrics as robust and reliable foundation for benchmarking GRN inference methods. The metrics capture context-dependent regulatory relationships, are sensitive to structural perturbations of GRN topology, and prioritize causal over purely correlational associations. In addition, they are robust to changing preprocessing choices such as normalization strategy and regression model selection.

### Simple models outperform complex algorithms in matched context

We used 11 metrics to benchmark GRN inference performance across 11 datasets. However, not all metrics were applicable to every dataset due to their specific requirements and underlying assumptions (Methods). For example, the Regression and Gene sets recovery metrics were applicable to all datasets, whereas TF recovery and WS distance metrics were only applicable to datasets with knockout perturbations (Extended Data Fig. 1-A, Extended Data Table 2). In addition to these basic requirements, we implemented a quality control procedure to ensure that the selected metrics provide a reliable basis for comparative evaluation (Methods). Specifically, for each metric–dataset combination, we applied three criteria: (i) Positive Control should perform better than Negative Control, (ii) at least one GRN model must exceed a minimum, metric-specific performance threshold, ensuring that the evaluation reflects biological signal rather than noise, and (iii) the variation in scores across GRN models must be sufficiently large to enable meaningful discrimination of model performance (Methods). Only metrics that satisfied these applicability criteria were retained per dataset (Extended Data Fig. 1-A). Overall, metric coverage ranged from Virtual cell (retained in 3 datasets) to Regression and Gene sets recovery (retained in all 11 datasets), with four metrics (Regression precision, Regression recall, Gene sets recovery, and TF binding) being applied in more than half of the datasets.

Our benchmark results show that performance depends strongly on both dataset and metric, with no single GRN inference method consistently ranking first across all dataset–metric combinations (Supplementary Fig. 1 and Supplementary Fig. 2). Overall, GRNBoost2, Pearson Correlation, Portia, and Scenic+ were the top-performing methods (excluding Positive Control; Fig. 3). The ranking was computed using a modality-balanced aggregation of normalized ranks across dataset–metric combinations (Methods). GRNBoost2, Pearson Correlation, Geneformer, Scenic, Portia, PPCOR, and scGPT were evaluated across the full set of 77 applicable combinations, whereas Scenic+, CellOracle, FigR, scGLUE, and GRaNIE were assessed on 11 combinations, as they were only applicable to multiomics datasets. scPRINT could not be applied to the Nakatake dataset due to its requirement for raw count data. Importantly, GRNBoost2 ranked first in 10 out of 11 datasets and second in the remaining one, demonstrating high consistency across datasets. GRNBoost2 also outperformed the Positive Control method, which is based on Pearson correlation computed using both inference and evaluation datasets. This indicates that the nonlinear regression models used in GRNBoost2 provide a stronger advantage than simply leveraging additional evaluation data in a correlationbased framework. Among transcriptomic GRN methods, GRNBoost2, Portia, and Pearson Correlation achieved the highest scores. Among the multiomics method, Scenic+ and CellOracle performed best.

**Figure 3:**
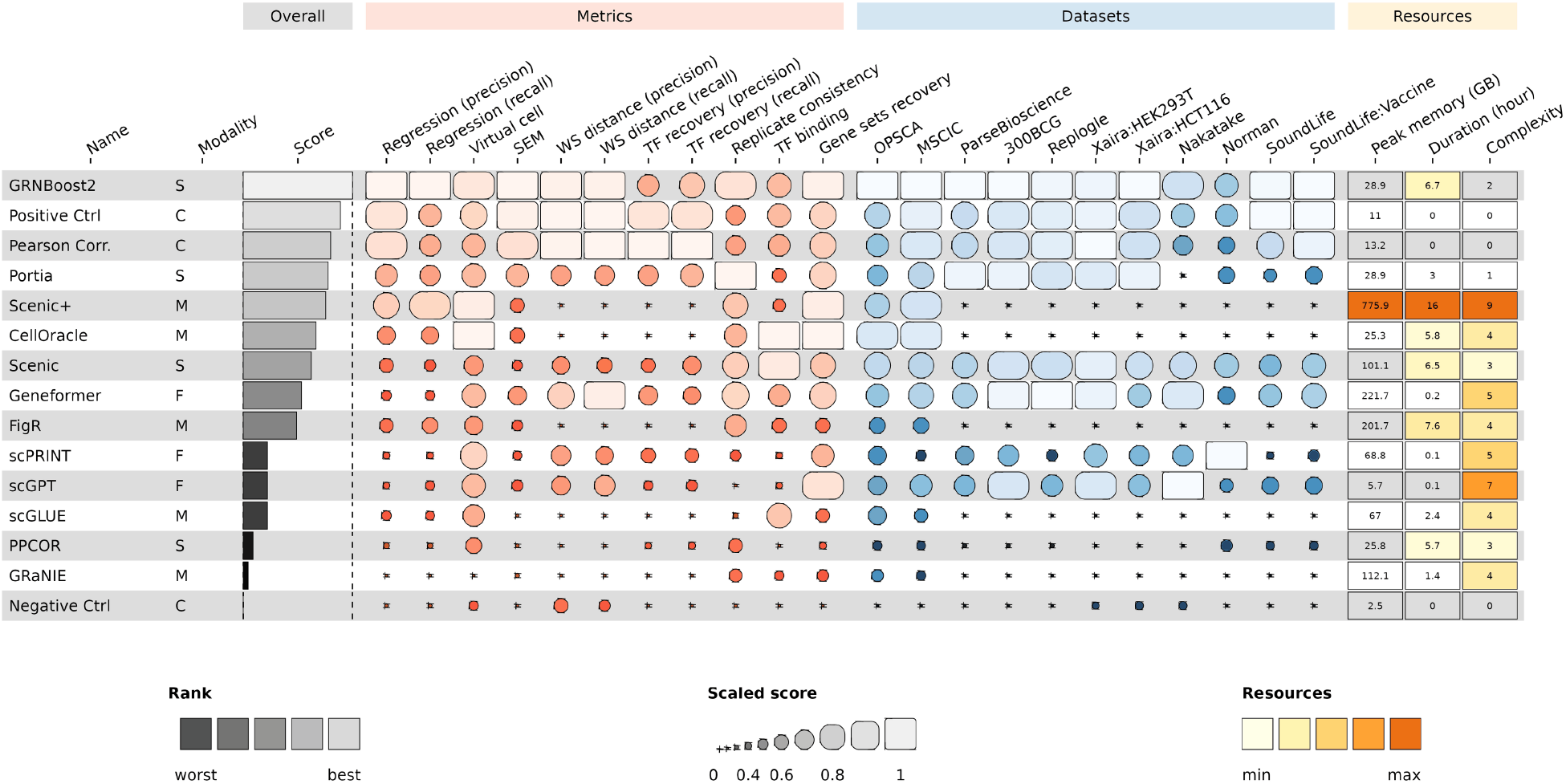
Overview of the GRN benchmark results. GRN inference methods are ranked based on an overall score computed in two steps: first, scores are min–max normalized within each dataset and metric combination; second, the median score was computed across all applicable dataset-metric combinations. Only metrics from datasets that passed quality control are included (Extended Data Table 2). Methods are categorized as: S (single-omics/transcriptomics), M (multiomics integrating scRNA-seq and scATAC-seq), C (control baselines), and F (foundation models). Asterisks (*) indicate cases where a specific dataset or metric is not applicable. Multiomics methods (M) were evaluated only on OPSCA and MSCIC datasets. Complexity scores reflect the subjective difficulty of method integration, considering factors such as documentation quality, dependency management, computational requirements, and ease-of-use.

Metric-specific evaluation showed that GRNBoost2 and Scenic+ performed best on regression-based metrics, followed by Pearson Correlation; Pearson Correlation led on WS distance, with GRNBoost2 and Geneformer ranking second and third; Pearson Correlation also led on TF recovery, followed by GRNBoost2 and Portia; GRNBoost2 achieved the highest SEM score, followed by Pearson Correlation and CellOracle; Portia and GRNBoost2 ranked highest for Replicate Consistency, with Scenic ranking third; CellOracle outperformed other methods on TF binding (followed by Scenic and scGLUE); GRNBoost2 outperformed others in Gene Set recovery followed by CellOracle and Scenic.

GRN modeling strategies among the methods integrated in geneRNIB broadly include (1) expression-based inference of TF–gene (or extended TF–peak–gene) interactions and (2) refinement using TF motif databases and chromatin accessibility. Different approaches are used to infer these interactions: co-expression (Pearson Correlation, GRaNIE, FigR), partial correlation/precision matrix (PPCOR, Portia), regression (GRNBoost2, Scenic, Scenic+, CellOracle), and foundation models (Geneformer, scGPT, scPRINT). Our results showed that regression-based methods, particularly GRNBoost2 and Scenic+ that use non-linear GBM, achieved higher performance than other approaches. Co-expression methods outperformed partial correlation/precision matrix methods (Supplementary Note 3). More notably, methods incorporating motif databases and chromatin accessibility performed lower than their expression-only counterparts: Scenic and Scenic+ performed below GRNBoost2, while GRaNIE and FigR ranked below Pearson Correlation. Foundation models showed intermediate performance, with Geneformer ranking highest among them, but remaining below coexpression baselines.

Together, our results show that simple, expression-based models, such as GRNBoost2 and Pearson Correlation, generally outperform more complex inference methods under our standardized, context-specific evaluation framework.

### Computational efficiency and ease of use of GRN inference methods

Beyond GRN inference performance, computational complexity and ease-of-use are important factors in method selection. We quantified computational complexity using CPU time and peak memory consumption. ease-of-use was assessed using a subjective score (1–10) for each method, based on integration complexity, including documentation quality, the number and severity of software dependencies, and the overall effort required to containerize these dependencies within a Docker environment for use in our reproducible platform. Methods such as Pearson Correlation and Portia required the lowest computational resources, completing runs in under 30 minutes and 1 hours, respectively, with peak memory below 30 GB (OPSCA dataset), and received the highest ease-of-use scores (10 and 9, respectively), reflecting their minimal installation overhead and straightforward interfaces. GRNBoost2, as the the top-performing methods, was among the methods with moderate complexity, requiring 6.7 hours and 28.9 GB of memory, with an ease-of-use score of 8. In contrast, deep learning-based methods such as Geneformer and scGPT, while computationally efficient in runtime (0.1–0.24 hours), scored substantially lower on ease-of-use (5 and 3, respectively). Scenic+ was the most demanding method across both dimensions: it consists of a multi-stage pipeline with numerous software and data dependencies (e.g., TF motif resources), a runtime of 16 hours, a peak memory footprint of 776 GB—an order of magnitude higher than most other methods—and the lowest ease-of-use. Overall, these results show that computational cost and ease-of-use vary substantially across methods, underscoring the need to consider these practical constraints alongside GRN inference accuracy when selecting an approach.

### Metric standardization is critical for GRN inference benchmarking

Any benchmarking study depends fundamentally on the fairness and robustness of the evaluation metrics. Systematic biases in these metrics can obscure true performance differences and lead to misleading conclusions. In our analysis, we identified two major sources of such bias: network topology and experimental variability. The inferred networks showed substantial variation in TF coverage, target gene coverage, and overall network density (Fig. 4-A and Supplementary Fig. 7-A and B). For example, GRNBoost2 exhibited broader TF and target coverage compared to Pearson Correlation, while maintaining lower network density. These structural differences can directly influence metric outcomes. For example, the Regression metric fits one model per gene using inferred TF as predictors. Therefore, the number of regulators alone, even if the inferred TF-gene connections are not accurate, can artificially improve performance. Experimental variability represents another confounding factor. For example, WS distance metric evaluates the accuracy of an inferred TF-gene edge by measuring the strength of the shift in target gene expression upon perturbation of the TF. Thus, a GRN centered around a TF with large global expression shift will score highly, even if the inferred edges are not accurate.

**Figure 4:**
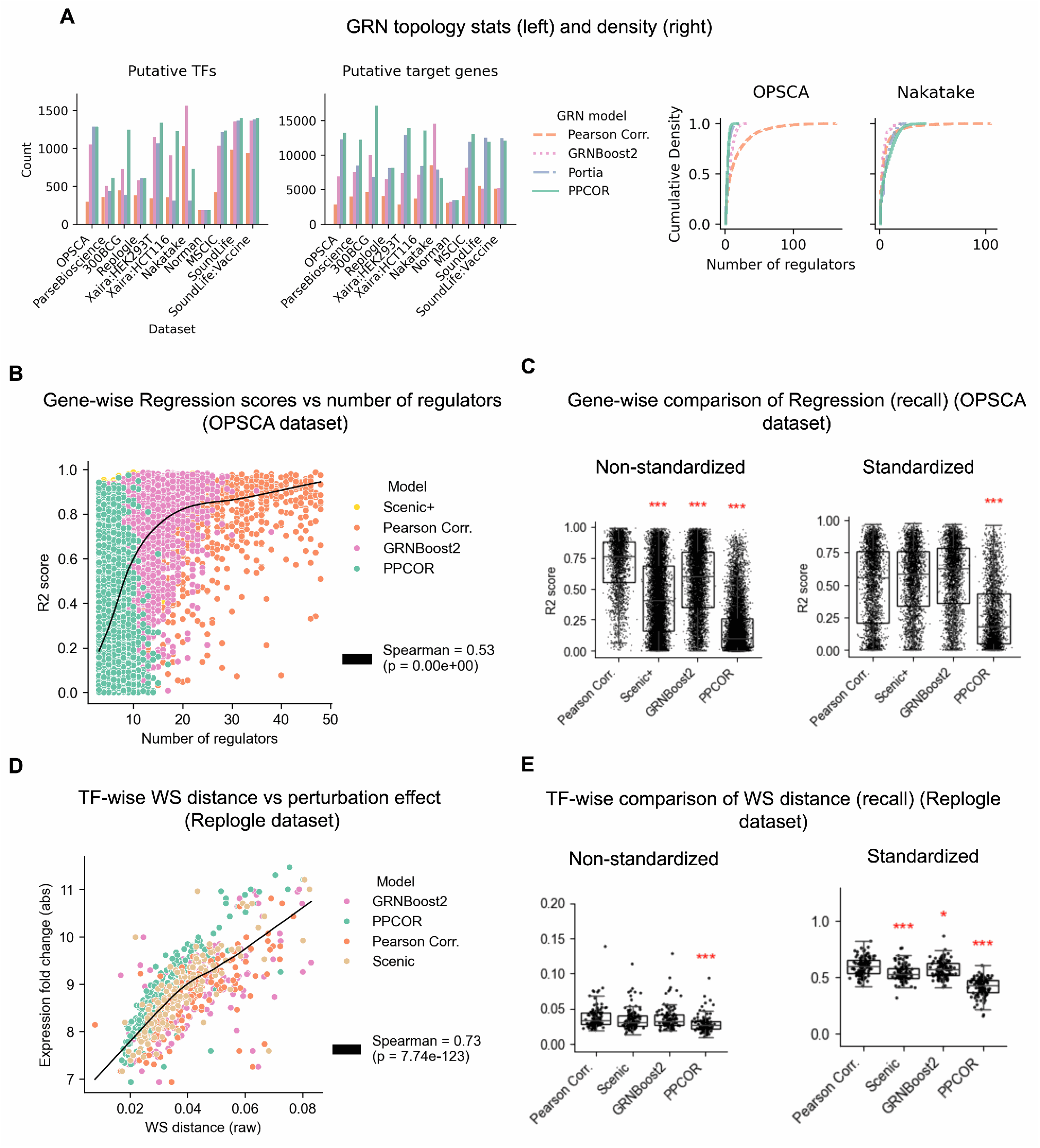
Analysis of metric standardization. **(A)** Topological statistics of GRN models for different datasets (left) and in-degree centrality calculated for selected methods (right). **(B)** Gene-wise performance of Regression metric as a function of the number of putative regulators. The curve represents a LOWESS (locally weighted scatterplot smoothing) fit. **(C)** Comparison of standardized versus non-standardized Regression metric. **(D)** Transcription factor (TF)-wise performance of the WS distance metric as a function of the perturbation strength, measured as log fold change of perturbed samples relative to control. **(E)** Comparison of standardized versus non-standardized WS distance. Red asterisks indicate the significance level (Mann-Whitney U test, FDR-corrected) when comparing Pearson Correlation model scores against other models (* *p <* 0.001, ** *p <* 0.01, * *p <* 0.05).

To address these issues, we standardized each metric by explicitly accounting for these confounding factors. For example, for Regression metric, we defined a consensus set of regulators per gene to control for variability in the number of predictors (Methods); for WS distance metric, scores were normalized relative to TF–specific background distributions to account for differences in perturbation strength (Methods). To quantify the impact of such standardization, we compared results obtained with and without these corrections for these two metrics, which have been used in previous studies without such adjustments [4, 8, 5]. For the Regression metric, gene-wise performance across models was positively associated with the number of regulators, indicating a potential confounding effect (Spearman *ρ*=0.54; *p <* 0.05; Fig. 4-B). Notably, comparative analysis showed that the non-standardized metric favored Pearson Correlation, which produces a denser network, leading it to significantly outperform all other methods, including Scenic+ and GRNBoost2, as confirmed by a rank test (Fig. 4-C). However, this advantage was no longer observed after standardization. Similarly, for the WS distance metric, TF-wise scores were strongly influenced by perturbation log fold changes (calculated relative to control), indicating a potential confounding effect (Fig. 4-D). In contrast to the Regression metric, Pearson Correlation significantly outperformed other methods only after standardization (*p <* 0.05, paired t-test; Fig. 4-E), whereas this advantage was not observed under the non-standardized metric.

Together, these results show the importance of our standardization toward a fair benchmark, allowing performance differences to more accurately reflect the accuracy of inferred regulatory relationships.

### Top GRN models show consistent performance across genes and TFs

To better understand what drives differences in GRN inference performance, we investigated whether high-performing models share common topological properties or infer regulatory elements such as TF and genes that tend to score higher. Topological overlap analysis, measured by Jaccard similarity, revealed no consistent pattern linking performance to shared network structure (Supplementary Fig. 5-A). For instance, GRNboost2 as a top-performing model showed higher edge-level overlap with Pearson Correlation and Scenic+ (other two competitive models), and higher TFs and target genes with PPCOR and Portia (less competitive models). This indicates that performance is not simply driven by converging on a common set of easy-to-score regulatory interactions. In addition, the overall low overlap of regulatory edges—even among top-performing models—indicates that different methods can capture distinct regulatory interactions while still performing comparably.

To further dissect this, we compared gene-wise and TF-wise performance scores derived from the Regression and WS distance metrics, respectively, across models. We calculated Spearman correlations between these scores and observed high cross-model agreement for gene-wise performance (0.57–0.89; Fig. 5-A), and relatively lower agreements for TF-wise scores (0–0.44; Extended Data Fig. 1-G), indicating that certain genes—and to a lesser extent TFs—are consistently better predicted across models. Importantly, top-performing models showed stronger agreement with each other than with less competitive methods. For example, GRNBoost2 exhibited correlations of 0.83 and 0.85 with Pearson Correlation and Scenic+, respectively, but only 0.57 with PPCOR for gene-wise scores (Fig. 5-A). Further visualization revealed that, despite an overall semi-linear relationship in gene- and TF-wise scores across models, top-performing models consistently achieve higher scores even for shared sets of genes and TFs (Fig. 5-B and Extended Data Fig. 1-F). For example, GRNBoost2 exhibited systematically superior gene- and TF-wise scores compared to PPCOR. Overall, these results show that top-performing models do not merely select easy-to-score regulatory edges, but result in more accurate relationships that is consistently reflected across a broad range of genes and TFs.

**Figure 5:**
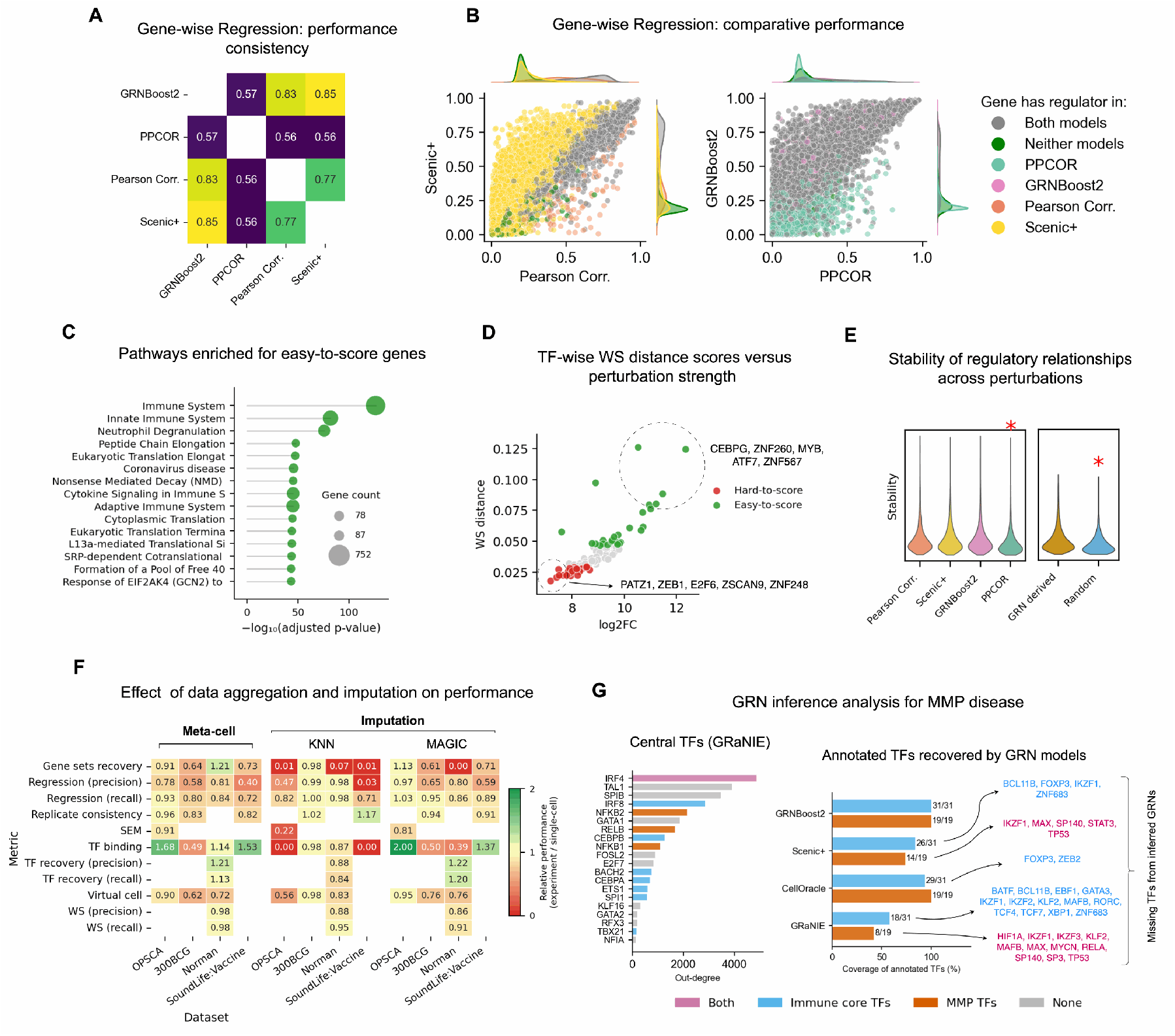
Performance analysis by gene and TF-wise evaluations, data aggregation and imputation, and MMP disease example. **(A)** Consistency of gene-wise Regression (precision) scores across models, measured by Spearman correlation. **(B)** Gene-wise Regression (precision) scores across different models. **(C)** Enrichment analysis of top-performing genes in the Regression metric. Adjusted *p*-values indicate the significance of enrichment, and gene count represents the number of genes contributing to each pathway. **(D)** TF-wise WS distance scores as a function of absolute log fold change (perturbed versus control) for top- and low-performing TFs. The top five TFs from each group are labeled. **(E)** Stability of regulatory relationships, defined as the mean-to-variance ratio of inferred TF–gene weights across perturbation subsets. The right panel compares GRN-derived regulators with randomly assigned TFs. Asterisks indicate statistically significant differences (*p <* 0.05). If not specified otherwise, the results are obtained for OPSCA dataset. **(F)** (left) Results of the data aggregation (meta-cell) analysis versus single cell data. Each value represents the relative performance of the aggregated data over the single cell data. The scores show the mean of two GRN inference models of Pearson Correlation and GRNBoost2 used for this experiment. Considering that multiple aggregation were performed, we used Leiden resolution of 9 (middle resolution) for this plot; (right) Performance evaluation for the imputed data (KNN and Magic) compared to the original single-cell data. The scores show the mean relative performance of imputed versus single cell data, calculated across two GRN models of Pearson Correlation and GRNBoost2 used for this experiment. **(G)** GRN inference in MMP disease: (left) GRaNIE highlights core immune markers and MMP-associated TFs as central regulators; (right) Number of recovered core immune markers and MMP-associated TFs across models; missing annotated TFs are indicated.

To further assess whether there exists a biologically meaningful subset of regulatory interactions that are consistently well- or poorly predicted across models, we systematically analyzed easy- and hard-to-score genes and TFs (Methods). Specifically, we identified the top and bottom 25% of genes and TFs based on gene-wise and TF-wise performance scores averaged across all models. For genes, using the OPSCA dataset composed of PBMCs, we obtained around 3,000 genes in each group (out of around 12,000), with mean R^2^ values of 0.70 and 0.02 for easy and hard genes, respectively. Easy genes showed 10× higher median expression, 44× greater variance, and were associated with 2.1× more regulators (Supplementary Fig. 5-C), suggesting their involvement in coordinated regulatory programs. Gene set enrichment analysis showed that easy genes were strongly enriched for core PBMC programs, including translation and ribosome biogenesis, cytokine production, inflammatory response, T cell receptor signaling, antigen receptor signaling, and neutrophil degranulation (Fig. 5-C). In contrast, hard genes showed overall weak enrichment (max *p*_*adj*_ ≈ 10^−3^ compared to *<* 10^−100^ in easy genes; Supplementary Fig. 5-B) with fewer genes per pathway (median 34 compared to 87 in easy genes), indicating that these genes are not concentrated within specific biological programs but are instead scattered across diverse processes. It included terms such as cilia assembly and cell cycle checkpoints, pathways that are not central to immune regulatory programs (Supplementary Fig. 5-B). For TFs, using Replogle dataset composed of K562, we identified 26 TFs in each group (out of 103), with mean WS distances of 0.065 and 0.026 for easy and hard TFs, respectively. Easy TFs were more central in inferred networks (69 vs. 48 edges on average) and induced 3-fold larger expression changes upon perturbation (Fig. 5-D). Easy-to-score TFs tended to include lineage-associated regulators with large regulons (e.g., MYB, CEBPG) as well as broad transcriptional repressors such as KRAB-domain zinc finger proteins, whose perturbation induced widespread transcriptional changes. In contrast, hard-to-score TFs were more often pleiotropic or context-dependent, including stress-responsive regulators (e.g., NFKB1, ATF3) that act as signal-dependent secondary responders and exhibit comparatively modest transcriptional shifts upon knockdown.

Overall, these results show that GRN inference performance is not uniform across genes and regulators, but is instead shaped by intrinsic biological properties. Genes that are highly expressed, variable, and embedded in coordinated regulatory programs—and TFs that induce strong and widespread perturbational effects—are consistently better predicted across models. This highlights that dataset-specific biological context fundamentally constrains what aspects of regulatory networks can be reliably inferred.

### Top GRN models produce regulatory interactions that are stable across perturbations and donors

GRN inference methods generally rely on gene co-variability to infer regulatory interactions; however, successful inference should leverage biologically meaningful variation rather than spurious associations arising from sampling effects or specific perturbation compositions. We analyzed this using five independent runs, where in each run 80% of the samples from the OPSCA perturbation dataset were randomly selected without replacement (Methods). For each GRN topology inferred by different models, we re-estimated regulatory weights using a per-gene regression approach while controlling for the number of regulators. As a baseline, we constructed model-specific random GRNs by assigning the same number of TFs to each gene. To quantify stability, we evaluated the consistency of inferred TF–gene weights across subsets, defined as the mean-to-variance ratio of weights across folds. We found that (1) GRN-derived regulators exhibited significantly more stable weights than randomly assigned TFs, and (2) top-performing models, including GRNBoost2, Scenic+, and Pearson Correlation, showed significantly higher stability compared to the lower-performing PPCOR model, as confirmed by rank-based tests (Fig. 5-E). We further repeated this analysis but this time using all perturbations while leaving one donor out (out of three donors) and calculating stability across donors. Consistent with the previous results, GRN-derived relationships remained more stable than random assignments, and top-performing models showed higher stability than less competitive models. These results suggest that top-performing GRN models capture regulatory structure that is more robust across perturbations and donors than random or lower-performing models, indicating that they recover relationships that are consistently supported by the data rather than those driven by specific sample compositions.

### geneRNIB dissects the impact of data imputation and aggregation on GRN inference performance

Many sophisticated GRN inference methods introduce simplifying assumptions that may introduce unintended biases. For instance, Portia estimates precision matrices under the assumption of a Gaussian distribution for gene expression, which may not accurately reflect the distributional properties of real biological data [26]. Similarly, methods such as GRaNIE and FigR employ data aggregation or smoothing strategies to alleviate the sparsity of single-cell data. To assess the impact of such aggregation strategies, we systematically evaluated the effect of pseudobulking on GRN inference performance (Methods). Specifically, we analyzed two aggregation schemes. First, we performed meta-cell aggregation by grouping cells based on unsupervised clustering (Leiden), generating increasingly coarse pseudobulk profiles across a range of clustering resolutions without using predefined labels (Methods).Second, we performed per-group aggregation by pseudobulking cells according to available biological annotations, including donor, cell type, perturbation condition, time point, and plate well.

Our results show that per-group aggregation consistently reduced GRN inference performance across datasets (Extended Data Fig. 1-H). However, meta-cell aggregation produced more complex effects depending on the dataset, evaluation metric, and the granularity of aggregation (Fig. 5-F and Supplementary Fig. 7-C). For example, aggregation generally reduced performance in the OPSCA, 300BCG, and SoundLife:Vaccine datasets but increased performance in the Norman dataset. At the metric level, the Virtual cell metric declined with increasing aggregation, whereas metrics such as TF binding showed a consistent improvement. Overall, these results indicate that while group-level pseudobulking tends to degrade GRN inference performance—consistent with previous studies reporting negative effects of data aggregation when evaluated with network-based metrics [27, 28]—this pattern does not uniformly apply to meta-cell aggregation.

Considering that data imputation is broadly used in single cell data analysis and also among GRN inference methods such as CellOracle, we evaluated its impact on GRN inference performance using two common imputation approaches: KNN (used in CellOracle) and MAGIC (Methods). We found that KNN imputation generally reduced performance across most metrics and datasets (Fig. 5-F). However, the effect of `MAGIC` was dataset- and metric-specific. For example, it reduced performance in the 300BCG and SoundLife:Vaccine datasets but generally increased performance in the OPSCA dataset. At the metric level, most metrics tended to decline following imputation, whereas the TF binding and TF recovery showed a general improvement.

Collectively, these results suggest that preprocessing strategies intended to improve signal-to-noise ratios, such as pseudobulking and imputation, do not universally enhance GRN inference performance.

### Top GRN models capture the biology of multiple myeloma precursor disease

To assess whether the performance measured by our metrics translates to real-world applications, we evaluated whether identified top-performing GRN models better capture the biology of multiple myeloma precursors (MMP), a group of plasma cell disorders—including Monoclonal Gammopathy of Undetermined Significance (MGUS) and Smoldering Multiple Myeloma (SMM)—that can progress to multiple myeloma. Specifically, we compared GRNs inferred by GRNBoost2 and PPCOR, representing top- and low-performing methods in our benchmark, for their ability to capture immune system biology and MMP-associated regulatory programs (Methods). The MMP dataset comprises two paired multiomics datasets (scRNA-seq and scATAC-seq) generated using 10x Genomics Multiome (10xGM) and DOGMA-seq (DOGMA). For GRN inference, both methods were applied to scRNA-seq data only. Network centrality analysis, defined as the number of target genes regulated by each TF, showed that GRNBoost2 preferentially ranked immune-related TFs among the most central nodes, whereas these TFs were largely absent from the central nodes identified by PPCOR (Supplementary Fig. 6). For example, GRNBoost2 identified marker TFs such as *ZEB2, BACH2, TCF4, SPI1*, and *EBF1* [29, 30, 31, 32] among the top hubs. In contrast, PPCOR prioritized TFs that are not well established in immune or MMP biology, such as *RFX6, NR2F1*, and *TFAP2B*. Consistently, GRNBoost2 preferentially targeted MMP-associated genes (128 genes extracted from the GWAS catalog; Methods), which exhibited significantly higher in-degree centrality—defined as the number of TFs regulating a gene—compared to other genes in the network across both 10xGM and DOGMA (Supplementary Fig. 6, *p <* 0.05). No such enrichment was observed for PPCOR. Together, these results indicate that GRNBoost2-derived networks more faithfully capture immune- and disease-relevant regulatory structure.

To better understand why GRNBoost2 outperforms multimodal approaches, we compared it with three methods—Scenic+, CellOracle, and GRaNIE—which showed moderate to competitive performance in our benchmark. Notably, these methods infer GRNs using paired scRNA-seq and scATAC-seq data, whereas GRN-Boost2 relies solely on scRNA-seq. All three methods prioritized curated immune marker TFs and MMP-associated TFs among central nodes (Supplementary Fig. 6). However, enrichment of GWAS-associated genes was less consistent than for GRNBoost2: for Scenic+, it was observed in 10xGM (*p <* 0.05) but not in DOGMA; for GRaNIE and CellOracle, it was observed in DOGMA (*p <* 0.05) but not in 10xGM (Supplementary Fig. 6). More importantly, all three multimodal methods failed to recover key TFs. Of 45 curated TFs, Scenic+, CellOracle, and GRaNIE missed 9, 2, and 21, respectively, across both DOGMA and 10xGM, whereas GRNBoost2 recovered all 45 (Fig. 5-G). Notably, CellOracle failed to recover *ZEB2*, a key regulator of immune cell identity and differentiation [29]; Scenic+ missed *IKZF1* (a pan-lymphoid master regulator and therapeutic target of lenalidomide in multiple myeloma [33]), key immune identity and differentiation factors including *BCL11B, ZNF683, BATF*, and *FOXP3* [34, 35, 36, 37], as well as MMP-associated TFs such as *STAT3, MAX*, and *SP140* [38, 39, 40]; Similarly, GRaNIE missed all TFs absent from Scenic+, and additionally key immune core TFs such as *EBF1, GATA3, RORC*, and *XBP1* [41, 42, 43]. Further analysis revealed that both TFs missing from CellOracle were also absent from the motif database used in this method (JASPAR2022; Methods). In contrast, 9 TFs missed by Scenic+ were all present in its motif database (cisTarget v10), indicating that they were likely lost during downstream chromatin enrichment scoring. Of the 21 TFs missed by GRaNIE, 11 were absent from its motif database (HOCOMOCO v12), while the remaining 10 were likely lost during peak–gene linking or regulatory correlation filtering.

Together, these findings showcase that improved performance on our evaluation metrics translates into a more complete and biologically meaningful recovery of context-relevant regulatory programs.

## Discussion and conclusion

GRN inference holds significant potential for elucidating cell behavior and phenotypes in both health and disease. Advancing this field requires a standardized benchmarking framework that can keep pace with rapidly evolving data types, inference strategies, and evaluation paradigms. Here, we presented geneRNIB, a living benchmark for GRN inference that enables continuous, context-specific, and standardized evaluation of GRN models across diverse perturbational and multiomics settings. By combining curated datasets, standardized metrics, and a dynamic leaderboard within a scalable computational infrastructure, geneRNIB provides a practical framework to help users select optimal methods and guide method development.

Our benchmark reveals a consistent trend: simple models based solely on gene expression, such as GRN-Boost2 and Pearson Correlation, generally outperform more complex approaches that integrate multiomics data or prior knowledge across multiple perturbation-based and predictive metrics (Fig. 3). This pattern is particularly evident across perturbation-informed and predictive metrics, including Regression, WS distance, TF recovery, and SEM. In contrast, multimodal and prior-informed methods tend to perform better on prior-based evaluations, such as TF binding, consistent with previous studies that relied on such benchmarks [10, 27, 3].Together, these results suggest that multimodal approaches preferentially capture global regulatory patterns rather than interactions tightly linked to the specific biological context, indicating a need to improve how prior information is incorporated to better model context-specific regulatory interactions. For example, many multiomics methods refine expression-derived GRNs using motif resources compiled across diverse tissues and conditions, which can filter out interactions that are active only in specific biological settings. This is illustrated by the comparison between Scenic/Scenic+ and GRNBoost2. Scenic and Scenic+ refine GRNBoost2-derived networks using TF motif and chromatin-accessibility priors; in our benchmark, this refinement reduced performance on context-specific metrics. Our MMP case study further showed that multiomics-based methods exclude important context-specific regulatory programs (Fig. 5-G), consistent with prior reports of missing known regulators for Scenic+ and CellOracle [3, 12]. Similarly, foundation models such as scPRINT, Geneformer, and scGPT, which are trained on heterogeneous datasets to learn general representations, show only modest performance in our benchmark, indicating that capturing broad regularities does not necessarily translate into accurate, condition-specific GRNs. Notably, simpler approaches also require substantially fewer computational resources and less implementation effort, further strengthening their practical advantages. Overall, these findings highlight a key unsolved challenge in GRN inference: balancing the use of global prior knowledge with the preservation of context-specific regulatory signals (Supplementary Note 4).

Aligned with our results, recent GRN benchmarking and perturbational studies also reported the limited improvement of more advanced algorithms over simple models. For example, CausalBench showed that networks derived from mean differences in target gene expression between observational and perturbational data performed competitively with sophisticated algorithms. In addition, Kernfeld et al. [44] showed that simply averaging gene expression values can outperform advanced algorithms in predicting perturbation effects. Extending these observations across a larger datasets and metrics compendium, we show that loss of context introduced by prior integration, along with simplifying assumptions such as data aggregation and imputation, can limit performance gain of more complex inference models.

Our results show that GRN inference should be evaluated within a defined biological context. The omics data used for inference are intrinsically linked to the underlying biological and experimental context, including tissue, cell type, disease state, and experimental condition. We show that context-dependent properties such as gene expression level and variability substantially influence inference performance (Supplementary Fig. 5). Consistent with this, GRNs inferred directly from our datasets outperformed networks derived from external resources on our context-sensitive evaluation metrics (Fig. 2-A). Even GRNs inferred from matched cell types under different experimental settings showed reduced performance, highlighting the importance of experiment-matched inference. These findings align with the broader understanding that regulatory interactions, including TF binding, chromatin accessibility, and co-regulatory relationships, are extensively rewired across cell types, physiological states, and experimental conditions [45, 2]. Experimental variation further compounds this effect through systematic shifts in omics profiles and batch effects [46]. Importantly, regulatory programs extend beyond transcriptional measurements alone. Cellular regulation operates across multiple molecular layers, including protein abundance, post-translational modifications, cofactor availability, and chromatin state, which cannot currently be captured within a single experimental assay [2]. Variation across these unmeasured layers likely contributes substantially to the context specificity observed even in transcriptomic GRN analyses. Restricting inference and evaluation to a well-defined biological setting therefore not only ensures that models are trained and assessed on coherent regulatory programs, but also provides a principled framework for partially accounting for the hidden molecular factors that shape regulatory wiring in each condition.

In geneRNIB, we considered multiple metrics for holistic assessment of GRN inference performance. Nonetheless, each metric embodies distinct assumptions and captures complementary dimensions of GRN quality, which can guide model selection toward particular applications. The SEM and Virtual cell metrics are naturally aligned with non-linear perturbation modeling and can guide GRN model selection toward interpretable frameworks for drug response prediction and perturbation planning [47, 48]. The Regression metric is valuable for applications requiring identification of functional upstream controllers—for instance, determining which TFs regulate disease-relevant genes. The TF recovery metric, which evaluates whether a GRN can correctly identify which TF was perturbed based on downstream transcriptional effects, is valuable in identifying the causal drivers of observed phenotypes. The WS distance metric, which focuses on functional TF–gene relationships at individual edge resolution, is informative for capturing regulatory interactions that are phenotypically important yet highly context-dependent. Gene set recovery evaluates whether a model correctly identifies TFs that regulate pathways active in a given biological context, making it valuable for applications that require pathway-level interpretability rather than precise edge-level reconstruction. Together, these metrics help guide the selection of GRN inference methods by matching model strengths to the specific requirements of different biological applications (Supplementary Note 4).

Our study has several limitations. First, with the exception of the TF binding metric—which directly measures physical TF–DNA interactions—the evaluation metrics do not distinguish between direct TF–gene regulation and indirect regulatory cascades, in which a TF regulates intermediate genes that subsequently affect the target gene. Incorporating additional experimental evidence will be important to refine the framework and enable more direct assessment of causal regulatory relationships (Supplementary Note 4). Second, current metrics do not adequately capture the sign or confidence of regulation (i.e., regulatory weight). In practice, these properties are either not explicitly considered or show limited sensitivity to perturbations (Supplementary Fig. 4). For example, regulatory weight is considered only in the Regression, TF binding, and Gene set recovery metrics, while regulatory sign is captured only by Virtual cell and SEM (Methods).Third, the predictive metrics employed in this study face challenges common to perturbation modeling, where dataset characteristics strongly influence model performance [49, 50]. The SEM and Virtual cell metrics, in particular, struggled with out-of-distribution predictions for genetic perturbations (Extended Data Fig. 1-A), which tend to produce more localized effects that do not propagate broadly across the transcriptome, making generalization more difficult. Similar trends have been reported previously, with strong performance on drug perturbation prediction [49] but limited success on genetic perturbations [44].Emerging approaches, such as STATE [51], may help improve performance in these more challenging settings. Fourth, we assumed that GRNs inferred under a given set of perturbations are transferable to unseen perturbations. While this is generally reasonable for targeted genetic perturbations, it may be less valid for broad-acting interventions such as drug treatments, which can induce perturbation-specific regulatory programs. Fifth, in multiomics settings, we assumed that chromatin accessibility remains constant between inference and evaluation due to limited availability of matched perturbation data. In reality, chromatin states are dynamic and responsive to perturbations. As more comprehensive multiomics perturbation datasets become available, incorporating these layers will improve benchmarking. Finally, current datasets are largely limited to cell lines and primary cells, with only the MMP dataset representing a primary tissue context. Expanding the inclusion of such datasets will enable more physiologically relevant benchmarking.

Overall, geneRNIB establishes a standardized and extensible benchmarking framework that can guide the development, evaluation, and application of GRN inference methods. We envision that this framework will advance the field toward more accurate and biologically meaningful models of gene regulation.

## Acknowledgments

This work was partially funded by Helmholtz-Zentrum Hereon. We acknowledge the support of the Maxwell computational resources operated at Deutsches Elektronen-Synchrotron (DESY), Hamburg, Germany, which facilitated parts of the computational analyzes. We extend our gratitude to OPSCA for providing the necessary computational infrastructure and to the staff of *Open Problems* for their invaluable assistance in troubleshooting and integrating geneRNIB into the OPSCA pipeline. M.S. was supported by a Joachim Herz Stiftung fellowship and the “Munich School for Data Science—MUDS”. Finally, we thank Pau Badia-i-Mompel, Artur Szalata, and Daria Romanovskaia for their insightful suggestions and constructive feedback and Anna Berezhna for her contributions to data visualization.

## Data availability

The multiple myeloma precursor (MMP), SoundLife, and MSCIC datasets were obtained from Gene Expression Omnibus under accession numbers GSE311602, GSE246714, and GSE194122, respectively. The inference and evaluation datasets together with all relevant data are hosted on s3://openproblems-data/resources/grn. Upon acceptance of this paper, we will deposit the data into a permanent repository.

## Code availability

The benchmark code is available at github.com/openproblems-bio/task grn inference. Additional code used to generate the figures and stability analyzes is hosted at github.com/janursa/genernbi supp. The entire benchmark code is provided under the MIT license. Upon acceptance of this paper, we will deposit the code into a permanent repository.

## Extended Data

**Extended Data Table 1:**
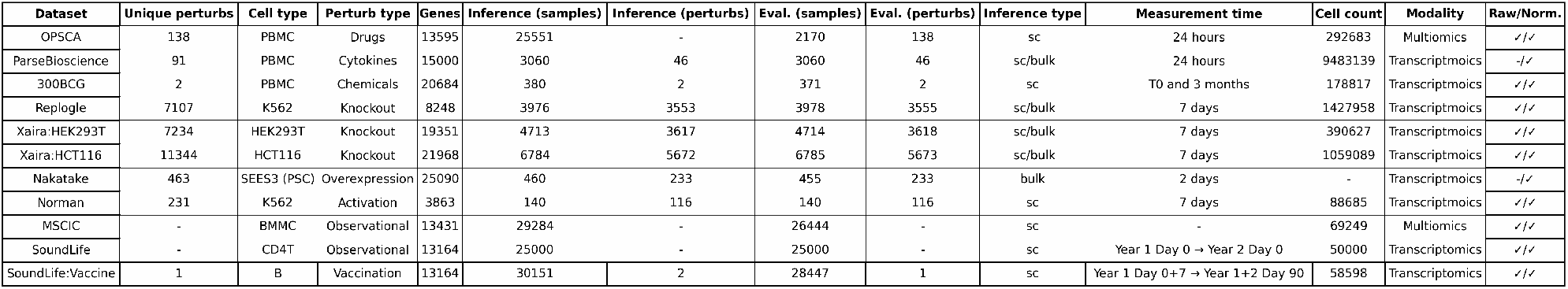
Overview of datasets in geneRNIB.

**Extended Data Table 2:** Overview of metrics and their applicability in geneRNIB. “Inputs” indicate the required files for the metrics. Applicable datasets are given for each metric, with * indicating if the metric passed the quality check for the given dataset. “Keep/total” indicate the ratio of datasets passing quality control checks. “Value (min/max)” reports the minimum and maximum values of the metric obtained across datasets. “Variability (min/max)” indicates the minimum and maximum of the metric variation across models measured per dataset, calculated as (max value - min value)/(mean value). A threshold of 0.2 was used. “Threshold” represents the global cut-off value defined for each metric. The performance of the Negative Control (NC) is reported for each metric, and the minimum of the global threshold and NC performance was used as the criterion in the quality control assessment.

| Metric | Summary | Inputs | Datasets | Keep/Total | Value (min/max) | Variability (min/max) | Threshold | NC |
| --- | --- | --- | --- | --- | --- | --- | --- | --- |
| Gene sets recovery | Evaluates GRNs by their ability to recover known gene sets | Inferred GRN, Evaluation data, Gene S | 300BCG*, MSCIC*, Nakatake*, Norman*, OPSCA*, ParseBioscience*, Replogle*, SoundLife*, SoundLife:Vaccine*, Xaira:HCT116*, Xaira:HEK293T* | 11/11 | 0.1/0.7 | 1.4/1.8 | 0.05 | 0.00/0.03 |
| Regression (precision) | Evaluates a GRN by the ability of TFs to predict target gene expression. It only evaluates top regulators. | Inferred GRN, Evaluation data, Known TFs list, Consensus regulators (Regression) | 300BCG*, MSCIC*, Nakatake*, Norman*, OPSCA*, ParseBioscience*, Replogle*, SoundLife*, SoundLife:Vaccine*, Xaira:HCT116*, Xaira:HEK293T* | 11/11 | 0.0/0.8 | 0.8/2.8 | 0.00 | 0.00/0.32 |
| Regression (recall) | Evaluates a GRN by the ability of TFs to predict target gene expression. It evaluates broader set of regulators. | Inferred GRN, Evaluation data, Known TFs list, Consensus regulators (Regression) | 300BCG*, MSCIC*, Nakatake*, Norman*, OPSCA*, ParseBioscience*, Replogle*, SoundLife*, SoundLife:Vaccine*, Xaira:HCT116*, Xaira:HEK293T* | 11/11 | 0.0/0.6 | 0.4/2.6 | 0.00 | 0.00/0.38 |
| Replicate consistency | Measures consistency of GRN predictions across biological replicates/groups. | Inferred GRN, Evaluation data | 300BCG*, MSCIC*, OPSCA*, ParseBioscience*, SoundLife*, SoundLife:Vaccine* | 6/6 | 0.5/0.8 | 1.0/2.1 | 0.30 | 0.05/0.32 |
| TF recovery (precision) | Measures ability to recover TFs using differential expression and GRN. It only evaluates available TFs in the GRN. | Inferred GRN, Differential expression data, Known TFs list | Norman*, Replogle*, Xaira:HCT116*, Xaira:HEK293T* | 4/4 | 5.6/9.5 | 1.2/5.3 | 2.00 | -0.66/1.40 |
| TF recovery (recall) | Measures ability to recover TFs using differential expression and GRN. It considers all TFs in the evaluation data. | Inferred GRN, Differential expression data, Known TFs list | Norman*, Replogle*, Xaira:HCT116*, Xaira:HEK293T* | 4/4 | 5.6/7.6 | 1.6/5.9 | 2.00 | -0.53/0.92 |
| WS (recall) | Evaluates a GRN by the shift in gene expression of target gene in response to TF perturbation. It evaluates broader set of TF-edge interactions. | Inferred GRN, Consensus edges (WS distance), WS distance background scores | Norman*, Replogle*, Xaira:HCT116*, Xaira:HEK293T* | 4/4 | 0.6/0.7 | 0.2/0.4 | 0.50 | 0.50/0.53 |
| WS (precision) | Evaluates a GRN by the shift in gene expression of target gene in response to TF perturbation. It evaluates top TF-edge interactions. | Inferred GRN, Consensus edges (WS distance), WS distance background scores | Norman*, Replogle*, Xaira:HCT116*, Xaira:HEK293T* | 4/4 | 0.7/0.8 | 0.3/0.5 | 0.50 | 0.51/0.54 |
| TF binding | Evaluates GRNs by their ability to match ChIP-seq TF binding data | Inferred GRN, Evaluation data, Ground Truth | 300BCG*, Norman*, OPSCA*, ParseBioscience*, Replogle*, SoundLife*, SoundLife:Vaccine*, Xaira:HCT116*, Xaira:HEK293T | 8/9 | 0.0/0.1 | 1.6/4.0 | 0.05 | -0.00/0.01 |
| SEM | Structural Equation Modeling for GRN evaluation. | Inferred GRN, Evaluation data, Known TFs list | OPSCA*, ParseBioscience*, Replogle*, Xaira:HCT116*, Xaira:HEK293T* | 5/6 | 0.0/0.2 | 1.7/3.0 | 0.01 | 0.00/0.01 |
| Virtual cell | Evaluate GRNs by their ability in predicting gene expression through neural networks-based virtual cell model. | Inferred GRN, Evaluation data | 300BCG*, Nakatake, Norman, OPSCA*, ParseBioscience*, Replogle, Xaira:HCT116, Xaira:HEK293T | 3/8 | 0.0/0.8 | 0.3/9.7 | 0.01 | 0.00/0.54 |

**Extended Data Figure 1:**
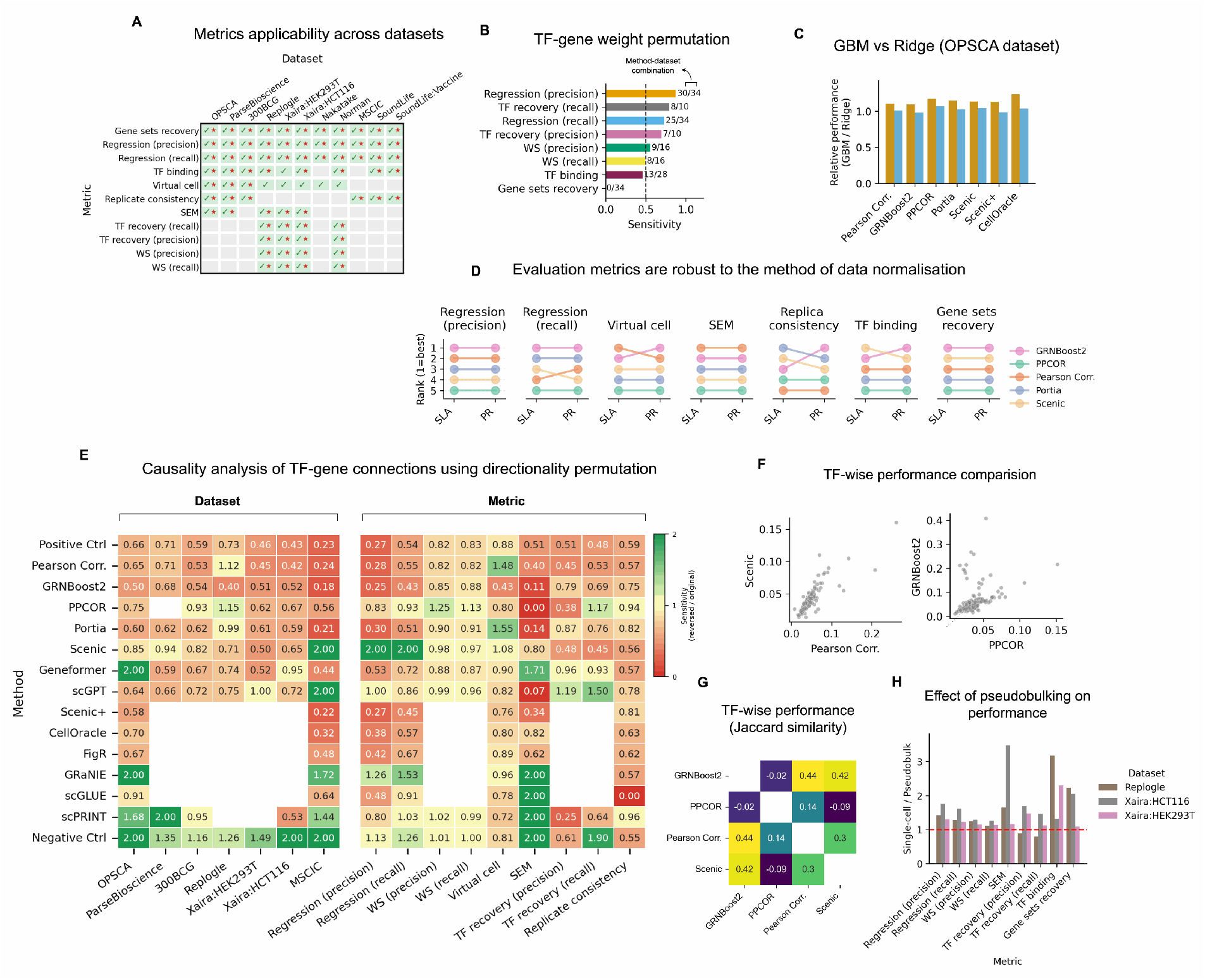
**(A)** Overview of metric applicability across different datasets. Green ticks indicate that a metric is applicable to the given dataset based on its requirements. The red star indicates whether the metric passed the quality control criteria for that dataset. **(B)** Results of permutation analysis of TF–gene weights. Sensitivity is defined as the performance of the original GRN relative to the permuted network. Numbers in front of the bars indicate the number of dataset–method combinations in which the original GRN outperformed the permuted version. **(C)** Performance comparison between Ridge regression and GBM for the Regression metric. **(D)** Performance comparison across two data normalization methods: shifted logarithmic approach (SLA) and Pearson residuals (PR). **(E)** Results of the causality analysis using TF–gene directional permutation. Each value represents the relative score of the permuted GRN compared to the original GRN (sensitivity). The color scale is capped at a maximum value of 2 for visualization purposes. Missing values indicate inapplicability of the metrics for given datasets. **(F)** The distribution of TF-wise scores WS distance scores. **(G)** Consistency of performance across models for TF-wise WS distance scores, captured by Spearman correlation. **(H)** Performance evaluation for the pseudobulked data compared to the single-cell data, calculated for GRN models inferred by Pearson Correlation.

## Methods

### Inference and evaluation datasets

The current release of geneRNIB includes 11 datasets (Extended Data Table 1), each identified by its corresponding tag. The OPSCA dataset originates from the 2023 Open Problems: Single Cell Perturbation competition [52]. It consists of single-cell drug perturbation data (10x Chromium) of 146 compounds together with multiomics measurements (scRNA-seq and scATAC-seq, 10x Multiome) of the baseline compound from the same experiment on PBMCs. The multiomics data was used for GRN inference data, while perturbation data was used as evaluation data. The MSCIC dataset originates from the NeurIPS 2021 Open Problems in Single-Cell Biology competition [16]. It is an observational dataset consisting of paired single-nucleus RNA-seq and ATAC-seq (10x Multiome) measurements from healthy bone marrow mononuclear cells (HBMMC) across 10 donors. We used multiomics data from 7 donors for GRN inference and RNA-seq data from the remaining 3 donors for evaluation. The 300BCG [53] dataset contains scRNA-seq measurements of chemical perturbations (LPS and RPMI) applied to immune cells (PBMC) from 38 donors, measured at two time points: baseline (0 months) and follow-up (3 months). We used baseline data for inference and the 3-month data for evaluation.

The SoundLife cohort was obtained from [54] and comprises approximately 12 million scRNA-seq PBMCs collected longitudinally from 96 donors using the 10x Chromium. The cohort includes healthy individuals, with measurements taken at baseline as well as following influenza vaccination across three consecutive years (years 1, 2, and 3) at multiple time points (day 0, 7, and 90). The dataset was provided with prior quality control and cell-type annotations. We selected approximately 100,000 single-cell profiles and divided them into four subsets: two for GRN inference and two for evaluation. The first pair follows a temporal split under baseline conditions (no vaccination), using the same donors across consecutive years, with GRNs inferred from CD4+ T cells at day 0 in year 1 (10 donors, randomly selected) and evaluated on CD4+ T cells at day 0 in year 2 from the same donors. The second pair also follows a temporal split but incorporates immune response dynamics, where GRNs were inferred from B cells at day 0 and day 7 in year 1 (10 donors, randomly selected) and evaluated on B cells at day 90 in both year 1 and year 2 from the same donors.

The Norman (K562) [55], Replogle (K562) [56, 57], Nakatake (SEES3 hESC) [58], ParseBioscience (PBMC) [59], and Xaira datasets (HCT116 and HEK293T) [60] are transcriptomic perturbation datasets covering perturbation types of gene activation, knockdown, overexpression, and cytokine stimulation, respectively. The Nakatake dataset was provided as bulk RNA-seq, while all others were single-cell. For all datasets, perturbations were randomly assigned to non-overlapping inference and evaluation subsets (unique perturbations per set). For large-scale single-cell datasets (Replogle, ParseBioscience, Xaira), we provide three versions for GRN inference: (i) the full single-cell dataset, (ii) a downsampled subset (restricted to TF perturbations for Replogle and Xaira, and to 10 out of 90 cytokines for ParseBioscience), and (iii) a pseudobulked version including all perturbations. All pseudobulking was performed by summing counts within the smallest grouping unit specific to each dataset, defined by combinations of perturbation, cell type, donor, and well. The pseudobulked datasets were used for all GRN inference methods except scPRINT, which requires single-cell counts, for which the full single-cell data was used.

Two datasets (Norman and Nakatake) were obtained from PEREGGRN [44], while the others were sourced directly from their original publications. All datasets were initially processed, quality-controlled, and normalized by their authors. We applied additional filtering to remove low-quality perturbations, cells, and genes. For OPSCA, details are provided in Supplementary Note 2. For other datasets, we required each gene to be detected in at least 10 cells and each cell to express at least 100 genes. For the large-scale knockdown datasets (Replogle, Xaira), we further ensured perturbation efficacy: for each perturbation, we compared expression against control using log2 fold change and t-test statistics, retaining only perturbations with log_2_ *FC <*− 0.54 and *p*_adj_ *<* 0.05. This reduced the Replogle dataset from 10k to 7k perturbations, the Xaira:HEK293T dataset to 7k perturbations, and the Xaira:HCT116 dataset to 11k perturbations (from 19k originally). The perturbation effects, calculated as the log fold change of individual perturbation versus control, is given in Supplementary Fig. 8-A.

### Evaluation metrics

We developed a comprehensive evaluation framework comprising multiple complementary metrics, each probing different biological aspects of network quality. In total, we implemented 8 main metric groups, which expand to 11 individual metrics when accounting for precision and recall variants explicitly reported for Regression, WS distance, and TF recovery. By default, we restricted all evaluations to TF-gene connections where the TF is included in a curated list of 1,638 known human TFs [61].

The WS distance was computed on single-cell data. The TF recovery metric was calculated on differential expression results obtained using the Wilcoxon rank-sum test between perturbed and control samples. All other metrics were computed on pseudobulked data, except for the observational data of MSCIC, where we used single cell data.

#### Regression

We constructed a regression model for each gene, using the expression profiles of its regulators as the feature space and the expression of the gene itself as the target. This method mirrors how several modern GRN inference techniques, such as GENIE3 [62], GRNBoost2 [19], and ENNET [63], build their regression models. We utilized the regulatory weights from the inferred GRNs to identify putative regulators for each gene and then extracted the expression data of these regulators (based on their gene correspondences) from the evaluation dataset to form the feature space. In this approach, the feature space for a given target gene varies between different GRN models. To standardize the number of features (that is, putative regulators) for a given gene across different GRNs, we introduced a variable, *m*_*j*_, which controls the number of regulators selected for each target gene *j*. For each gene, we ranked the putative regulators according to their inferred weights and selected the top *m*_*j*_. For GRNs that include negative regulations, we took the absolute values of the weights. To handle cases where multiple candidate regulators had the same rank, we injected arbitrarily small random values into the network to break ties and ensure a unique ranking.

For each gene *j*, we restricted *m*_*j*_ to be identical across all GRNs to ensure fairness. To standardize the calculation of *m*_*j*_, we first identified the number of inferred regulators of gene *j* per GRN model. Let 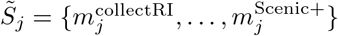 be the actual counts of regulators of gene *j* for all GRNs. Intuitively, choosing 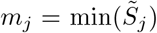 would privilege methods that produce accurate predictions among their very top regulatory edges. On the other hand, choosing 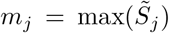 would penalize GRNs with lower coverage (that is, sparser networks). We generalized these ideas and instead computed 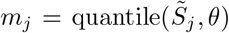 as the quantile function across all GRNs with cut-off *θ*. This approach covers the cases 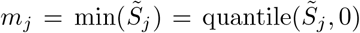 and 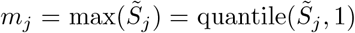. We used the `quantile` function of the `NumPy` Python package, which performs linear interpolation. For cases where the number of putative TFs is less than *m*_*j*_, we randomly selected the remaining TFs to fill the gap. Considering that the ground truth number of regulators for a given target gene is unknown, the choice of *θ* can serve as a trade-off between false positive and false negative rates.

The number of consensus regulators for these *θ* cut-offs are given in Supplementary Fig. 8-B. We selected two cut-off values of 0.25, and 0.75, representing the low and high number of allowed regulators. The scores resulting from these cutoff values are referred to as *Regression*precisionand *Regression*recall, respectively. We observe that the number of regulators vary from 0 to 20 for for *Regression*precision, where the most relevant regulatory interactions are captured, while *Regression*recallcovers a large space of regulators. The number of genes with at least one regulator is shown in Supplementary Fig. 8-D. This number varies from a few thousands to *Regression*precisionto tens of thousands to *Regression*recall. In the calculation of final scores, we skipped those genes that did not have at least one regulator shared across the models, as they would inflate the number of random predictions due to missing genes.

We used a 5-fold cross-validation to evaluate the feature space obtained from the GRNs. The scores were averaged across the genes to obtain the final score. We used Ridge as default model.

#### Structural equation models (SEM)

SEMs are causal models that describe how a gene’s steady-state is determined directly (by its regulators) and indirectly (by exogenous inputs). SEM has been previously employed in GRN inference [64, 65, 66, 67, 68]. here, we use a similar concept for the evaluation of inferred GRNs.

Let *x* ∈ ℝ^*m*^ be a preprocessed gene expression profile, and *u* a vector of exogenous variables (e.g., compound effects). The steady state of the system can be described by the following linear SEM:

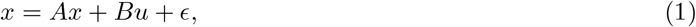

where *A* ∈ ℝ^*m*×*m*^ is an endogenous operator (a GRN) representing the direct effect of a regulator on its target genes, *B* ∈ ℝ^*m*×*d*^is an (exogenous) actuation map representing the direct drug effects on gene activity, and *ϵ* is an unexplained noise vector. Assuming *I* − *A* is invertible, then the steady state has a closed form:

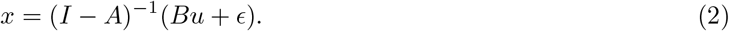

(*I* − *A*)^−1^ can be interpreted as an operator that sums all direct and indirect regulations between genes. Indeed, this is visible from the Neumann series:

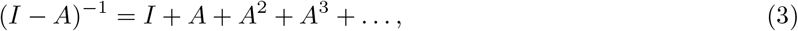

where higher degrees correspond to long-range propagation.

Now, we define Δ*x* as the difference between steady states after administration of a drug with effect Δ*u*, that is,

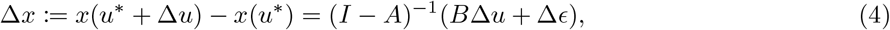

where *u*^∗^ is the steady-state value of *u* in a matched control. For notational convenience, we also define the exogenous shock *δ* = *B*Δ*u*. Perturbation Δ*x* now becomes:

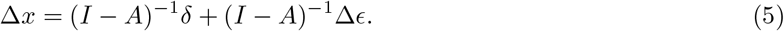

Under exogeneity assumption (i.e., the absence of correlation between the independent variables and the error term), 𝔼 [Δ*ϵ*|Δ*u*] = 0. Therefore, the expected value of a perturbation Δ*x* is given by:

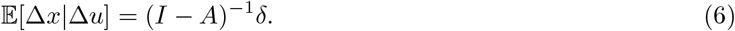

In causal terminology, the SEM predicts the do-effect of *u*, denoted by 𝔼 [Δ*x*|Δ*u*].

Given an inferred GRN *A*, we aim to evaluate its performance as a linear SEM. We measure Δ*x* as the difference between treated cells and matched controls. we defined matching in different granularity depending on the datasets, with matching covariates of cell-type, donor, plate name, and well.

However, quantifying the exogenous shock *δ* is less trivial. Since the exact effect of each compound on each gene is unavailable, we include the estimation of *δ* as part of the problem. This is achievable by splitting the genes into two sets: a set of *reporter genes* and a set of *evaluation genes*. We define reporter genes as the genes that are likely to be directly influenced by interventions. We included all TFs in the reporter gene set. Indeed, since pharmacology acts proximal to the network (e.g., TFs, kinases), the expression of downstream genes is altered mostly by propagation through the network. We also included immediate early genes in the reporter gene set. The list was obtained using the `scCustomize (v3.2.0)` R package [69]. All the remaining genes were considered for the evaluation. Also, we split the perturbation profiles into 3 sets to avoid data leakage: controls, training set and validation set. By splitting the genes and samples into respectively 2 and 3 sets, we ended up with 6 sets, which were grouped into 4 sets as illustrated in Supplementary Fig. 8-F.

In summary, our evaluation approach is as follows:

- Extract a mask *M* ∈ {0, 1}^*m*×*m*^ from the inferred GRN *A*, where *M*_*ij*_ = 1 when *A*_*ij*_ ≠0.
- For each column *j*, fit one ElasticNet regression model on the controls, only including the features for which the corresponding elements in *M*_·*j*_ are non-zero. Build a new matrix *A* from the linear coefficients, which will serve as an initial estimate for the endogenous operator in the SEM.
- Solve the minimization problem min 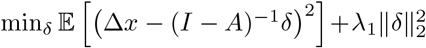 on both training and validation sets (reporter genes only) to estimate the shocks *δ*. Since *A* is constant, a closed-form solution exists.
- Solve the minimization problem min 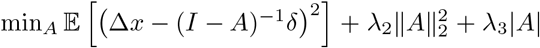 on the training set (evaluation genes only) to infer *A*.
- For each evaluation gene *j*, compute the Spearman *ρ* between Δ*x*_*j*_ and ((*I* − *αA*)^−1^*δ)*_*j*_, using only the perturbation profiles from the validation set. Average the correlation coefficients. To balance network fidelity with gene-level coverage, we run the entire evaluation procedure twice: once using only the genes present in the inferred GRN, and once using a set of 3,000 highly variable genes (HVGs). HVGs are identified using `Scanpy` with the Seurat flavor. The final score is defined as the average of the two results. This strategy balances inference precision—capturing what is retained in the GRN model—with coverage of genes that are important for the cell type, which serves as a surrogate for recall, ensuring that both aspects contribute to the final evaluation.

#### Virtual cell

This metric is inspired by recent advancement in virtual cell (VC) and follows similar modeling approach to modern approaches such as State [70]. VC evaluates the utility of an inferred GRN for predicting cellular responses to perturbations by training a neural network that incorporates the GRN structure to predict perturbed gene expression from control expression. The underlying hypothesis is that a high-quality GRN should encode regulatory relationships that generalize to predict how cells respond to unseen perturbations.

We formulate this as a supervised learning problem where the model predicts the expression profile of a perturbed cell given (i) the expression profile of its matched control cell, and (ii) the identity of the perturbation. The model architecture integrates the inferred GRN through specialized layers that transform expression data according to the network topology. Specifically, for a GRN represented as an adjacency matrix *A* ∈ ℝ^*m*×*m*^, where *A*_*ij*_ denotes the regulatory effect of gene *i* on gene *j*, the model applies GRN-informed transformations at both the input and output stages (layers). The input transformation applies the GRN directly to the control expression: *x*_transformed_ = *x A*^*T*^, where *x* is the control expression profile. The middle layers encode perturbation information through learned embeddings combined with the transformed control expression via feedforward neural networks. The output transformation applies an inverse GRN operation: *y*_predicted_ = (*I* − *αA*^*T*^)^−1^*y*_decoded_, where *α* = 0.1 is a dampening factor and *y*_decoded_ is the output from the middle layers. This inverse transformation captures both direct and indirect regulatory effects through the Neumann series expansion (*I − αA*^*T*^)^−1^ = *I* + *αA*^*T*^ + (*αA*^*T*^)^2^ + · · ·, analogous to the SEM formulation.

The encoder and decoder parts of the neural architecture contain 2 layers each with parametric ReLU activation functions. The model was trained with the Adam optimizer, using a batch size of 512, a learning rate of 10^−3^ and a weight decay of 10^−5^.

We evaluate prediction quality using 5-fold cross-validation stratified by perturbation identity to ensure the model generalizes to unseen perturbations. For each fold, we train the model on one subset of perturbations and evaluate on held-out perturbations. The matching between perturbed samples and control samples is performed based on dataset-specific covariates (e.g., cell type, donor, plate, well) to account for technical and biological confounders. Model training optimizes the mean squared error between predicted and true perturbed expression profiles using the Adam optimizer with learning rate scheduling.

The metric is computed as the coefficient of determination (*R*^2^) averaged across all genes and folds:

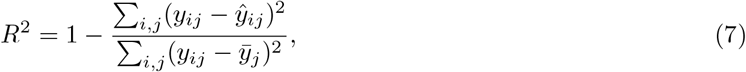

where *y*_*ij*_ is the true expression of gene *j* in sample *i, ŷ*_*ij*_ is the predicted expression, and 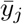 is the mean expression of gene *j* across samples. We compute this metric separately for two gene sets: (i) the top 1,000 most connected genes in the GRN, emphasizing genes central to the network structure, and (ii) the top 3000 highly variable genes, focusing on genes with the most dynamic expression patterns. The final virtual cell score is reported as the average of these two metrics. This dual evaluation ensures the metric captures both network-centric and expression-centric predictive performance.

#### Wasserstein (WS) distance

As an alternative evaluation metric, we adopted the WS distance from [8]. In this metric, for a given edge representing a pair of TF-target gene, we calculated the WS distance between the distribution of the target gene’s expression in observational (control) data and its expression in interventional data of the corresponding TF. A true edge is expected to yield a higher distance. The final score for a GRN was obtained by averaging the WS distances across all edges. To standardize comparisons between different models, which may vary in their coverage of TFs and target genes, we implemented two key measures. First, for each TF given in the interventional data, we calculated a background distribution of WS distances by randomly selecting 1,000 edges. Subsequently, instead of using raw WS distances to evaluate an edge, we computed the percentile rank of the edge within its respective background distribution. This standardization accounts for potential technical variability in the gene expression of two different interventions. Second, to standardize the number of edges considered for each TF, we adopted a consensus approach similar to previous section with two threshold values for *θ* (0.25 and 0.75). A *θ* of 0.25 emphasizes precision, whereas a *θ* of 0.75 emphasizes recall. The distributions of edge counts for each *θ* threshold are shown in Supplementary Fig. 8-C. If a network contains fewer targets for a given TF than expected by the evaluation, we assign random targets (with replacement) from the background gene set.

#### Transcriptional factor (TF) recovery

This metric assesses the ability of a GRN to correctly identify the perturbed TF in a given experiment. First, we calculated TF activity scores for the TFs using the GRN topology and the corresponding gene expression data via the `decoupler` univariate linear model [71]. Only TFs with at least three target genes were considered. Considering that these activity scores can be confounded by inter-experiment variability, we computed a baseline score for each TF and use the relative improvement of the actual score over the baseline as the final evaluation score. For the baseline GRN, each TF was assigned to a set of randomly selected *n* target genes, where *n* equals the number of actual targets of that TF. We then performed a paired *t*-test comparing the actual TF activity scores to their baseline counterparts across TFs, and use the resulting *t* statistic as the evaluation metric. We refer to this metric as TF recovery precision. To further account for the comprehensiveness of the inferred network, we computed an additional metric called TF recovery recall. The key difference between these two metrics is that TF recovery precision includes only the TFs present in the inferred GRN, whereas TF recovery recall includes all TFs from the evaluation dataset. For TFs missing in the GRN, we assigned the median baseline score. This approach penalizes networks that are largely incomplete. For TF activity estimation, we used differential expression (DE) data derived from large-scale perturbation datasets (e.g., Replogle, Xaira: HEK293T, Xaira: HCT116). For each dataset, we compared gene expression between perturbed and control samples. DE analysis was performed using the `Scanpy rank_genes_groups` function with the *Wilcoxon rank-sum test*, and the resulting test statistic was used as the DE value for downstream TF activity calculation.

#### Replicate consistency

This metric evaluates the biological reproducibility of TF activity estimates derived from an inferred GRN by measuring their consistency across technical or biological replicates. The underlying hypothesis is that a high-quality GRN should produce TF activity scores that are stable across replicate samples from the same experimental condition, as true regulatory signals should be reproducible while noise-driven predictions would vary randomly.

For a given evaluation dataset containing replicate samples grouped by experimental conditions (e.g., perturbation × cell type combinations across different donors or batches), we first compute TF activity scores using the ULMP from `decouplerDr` [71]. For each TF *f* with target gene set 𝒯_*f*_ defined by the inferred GRN, ULM estimates activity scores across all samples through weighted linear regression of gene expression on the TF-target network topology. Only TFs with at least 5 target genes are considered to ensure reliable activity estimation.

To quantify consistency across replicates, we measure the dispersion of TF activity scores within each experimental group. For each TF *f* and group *g* (representing a unique perturbation × cell type combination), we compute the Mean Absolute Deviation (MAD) from the median:

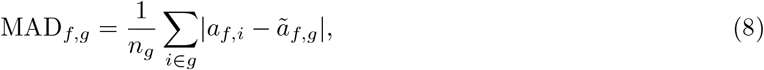

where *a*_*f,i*_ is the activity score of TF *f* in sample *i, ã*_*f,g*_ is the median activity across all samples in group *g*, and *n*_*g*_ is the number of samples in the group. We use MAD instead of coefficient of variation because it is more robust for small sample sizes (2-3 replicates per group) and does not require positive values. The overall dispersion for TF *f* is then computed as the mean of MAD_*f,g*_ across all groups.

We convert dispersion to a consistency score by normalizing and inverting: consistency increases as dispersion decreases. Let MAD_*f*_ denote the mean dispersion of TF *f* across all groups. We normalize dispersions using the 95th percentile as an upper bound to avoid sensitivity to outliers, then compute consistency as:

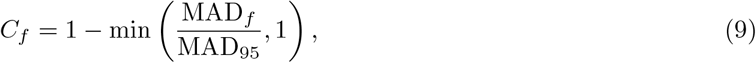

where MAD_95_ is the 95th percentile of all TF dispersions. This yields *C*_*f*_ ∈ [0, 1], where *C*_*f*_ = 1 indicates perfect consistency (no variation across replicates) and *C*_*f*_ = 0 indicates high dispersion.

To account for varying network sizes and densities across different GRN inference methods, we evaluate consistency across multiple network configurations. We systematically vary the number of top TFs included (ranked by degree centrality) from 10 to 300, evenly spaced across 10 configurations. For each configuration with *K* TFs, we select the top *K* TFs by their number of target genes, compute consistency scores for each TF, and calculate the mean consistency across the top *K* TFs (padding with zeros if fewer than *K* TFs meet the minimum target threshold). The final Replicate consistency score is reported as the mean across all 10 configurations. This averaging approach ensures that methods are evaluated fairly regardless of whether they produce dense or sparse networks.

#### Transcriptional factor (TF) binding

The TF binding evaluation metric quantifies the fidelity of predicted gene regulatory networks by comparing them against experimentally derived TF binding data from ChIP-seq–based resources. TF binding peaks were obtained through the GRETA pipeline [4], which integrates data from three complementary databases: ReMap2022 [72], ChIP-Atlas [73], and UniBind [74]. We collected TF binding data for four cell types: PBMC, K562, HEK293T, and HCT116. When cell type–specific data was unavailable, we included binding data from closely related tissues or cell types. Specifically, for PBMCs, we used data from immune-related cells and tissues; for K562 and HEK293T, we used data directly reported for these cell lines; and for HCT116, we included data reported under related colorectal cancer cell lines such as DLD-1, HT-29, and CACO-2. To construct TF–target ground truth networks, we first retrieved transcription start site (TSS) annotations from [75]. For each TF, binding peaks were intersected with promoter regions defined as ±1,000 base pairs upstream and 100 base pairs downstream of the TSS, accounting for strand orientation. This process yielded three ground truth networks per cell type—one for each source database. Summary statistics for the number of TFs and TF–target interactions are shown in Supplementary Fig. 8-E. We observed broad coverage for K562 and HEK293T, with 50–400 TFs depending on the source, and moderate coverage for PBMC and HCT116, with approximately 10–50 TFs per source. Supplementary Fig. 7-D shows the overlap of TFs and edges between the three sources. While the overlap in TFs was moderate to high (0.1–0.7), the overlap in TF–target edges was comparatively low (0.06–0.15).

Using these ground truth networks, we computed TF binding–based evaluation scores for each dataset and ground truth source. Considering the multiple datasets corresponded to the same cell type, we used the same ground truth network for evaluation. Specifically, the OPSCA, 300BCG, ParseBioscience, and SoundLife datasets were all evaluated using the PBMC ground truth, and the Replogle and Norman datasets were evaluated using the K562 ground truth. In total, we evaluated nine out of 11 datasets for which ground truth data was available.

To compute the TF binding metric, we compared the predicted TF–target interactions from the inferred GRN to the ground truth networks. For each TF present in the ground truth data, we extract the set of known target genes 𝒯_true_ from the binding database. Let *k* = |𝒯_true_| denote the number of true target genes for a given TF, and let *n* denote the total number of genes in the evaluation dataset. From the inferred GRN, we rank all predicted target genes of the TF by their absolute regulatory weights and select the top *k* predictions to form the set 𝒯_pred_. The precision for a given TF is calculated as the proportion of true targets recovered among the top *k* predictions: |𝒯_true_ ∩ 𝒯_pred_ |*/k*. To account for random chance, we normalize this precision by subtracting the expected precision under random selection and scaling by the maximum achievable precision. The expected precision by random chance is *k/n*, while the maximum precision is 1.0 (when all top *k* predictions are correct). The normalized precision for TF *f* is then computed as:

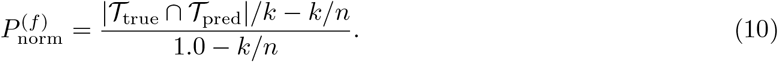

This normalization yields a score where 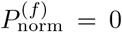 indicates random performance, 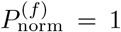 indicates perfect recovery, and negative values indicate performance worse than random.

We compute two aggregate metrics to capture different aspects of network quality. The *TF binding precision* score is calculated as the mean normalized precision across all TFs that are present in the inferred GRN:

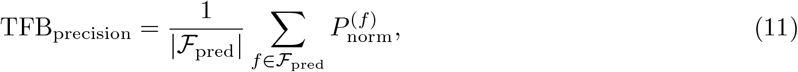

where ℱ_pred_ denotes the set of TFs in the ground truth that are also present in the inferred GRN. This metric emphasizes the quality of predictions for TFs that the method chooses to include.

To penalize networks with low TF coverage, we also compute the *TF binding recall* score, which averages the normalized precision across all TFs in the ground truth, assigning a score of zero to TFs missing from the inferred network:

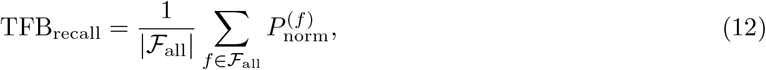

where ℱ_all_ is the set of all TFs in the ground truth, and 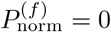 for TFs not present in the inferred GRN.

Finally, we combine precision and recall into a single F1 score:

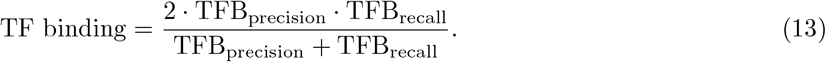

Since multiple ground truth databases are available, we compute these metrics separately for each database and report a weighted average, where the weight for each database is proportional to the number of TFs it contains.

#### Gene sets recovery

This metric evaluates whether the inferred GRN captures biologically meaningful functional relationships by assessing the enrichment of canonical gene sets among the target genes of TFs. The underlying hypothesis is that TFs regulating genes within the same biological pathway should exhibit significant enrichment for that pathway’s gene set. We assess this by comparing pathway activity in the evaluation dataset against pathway enrichment patterns derived from the inferred GRN. This metric is adopted from GRETA [4] with additional adjustments such as standardization of network topology and broader gene sets databases.

We first identify *active pathways* in the evaluation dataset using the ULM from `decoupler` [71]. For a given pathway *p*, we construct a network where the pathway name serves as the source and each gene in the pathway as a target with uniform weight. ULM then estimates pathway activity scores across all samples in the evaluation dataset. We determine pathway activity through a one-sample *t*-test, testing whether the mean activity score is significantly greater than zero. A pathway is considered active if *t*-test *p*-value *<* 0.05 and mean activity *>* 0. Next, we determine *enriched pathways* in the inferred GRN using Fisher’s exact test for each TF-pathway pair. For a given TF *f* with target gene set 𝒯_*f*_ and pathway *p* with gene set 𝒢_*p*_, we construct a 2 × 2 contingency table and perform a one-sided Fisher’s exact test to assess enrichment. To standardize for varying network densities across different GRN inference methods, we limit each TF to its top *K* targets ranked by absolute edge weight, where we set *K* = 100. A pathway is marked as enriched if at least one TF shows significant enrichment after Bonferroni correction for multiple testing across all tested pathways.

We evaluate GRN quality by comparing pathway activity (from the evaluation data) against pathway enrichment (from the inferred network). For each pathway, we classify it into one of four categories:

- True Positive (TP): Pathway is both enriched in the GRN and active in the dataset
- False Positive (FP): Pathway is enriched in the GRN but not active in the dataset
- False Negative (FN): Pathway is active in the dataset but not enriched in the GRN
- True Negative (TN): Pathway is neither enriched nor active

From these counts, we compute the gene set recovery precision, recall, and final score which is F1 score:

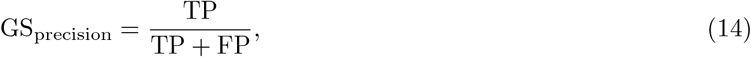

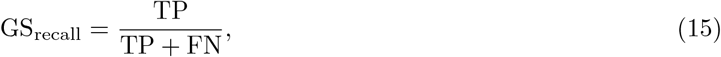

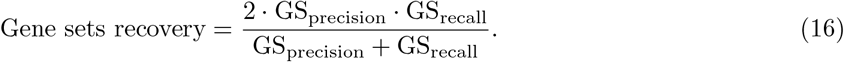

The precision metric measures how well the enriched pathways in the GRN correspond to biologically active processes, while recall captures the GRN’s ability to recover all active pathways. We evaluate gene sets from multiple databases including `MSigDB Hallmark 2020` [76], `KEGG 2021` [77], `Reactome 2022` [78], `GO Biological Process 2023` [79], `BioPlanet 2019` [80], and `WikiPathways 2019` [81]. The final scores are reported as the mean across all gene set collections. Pathways are filtered to include only those with 5 to 500 genes to focus on well-defined biological processes while excluding overly broad or narrow categories.

### Analysis of metric applicability across datasets

Each metric requires specific data properties, which limits its applicability across datasets. A summary of metric applicability is provided below.

#### Regression (precision / recall

Requires a sufficient number of samples to train a model that predicts target gene expression from TF gene expression. Applicable to all datasets.

#### WS distance (precision / recall

Requires single-cell data with TF genetic perturbations (KO/KD/overexpression). Applicable to the Replogle, Norman, and Xaira datasets.

#### TF recovery (precision / recall)

Requires TF perturbation experiments with sufficient samples to compute differential expression values. Applicable to the Replogle and Xaira datasets. Nakatake was excluded because it contains only five control samples, limiting differential expression analysis.

#### Replicate consistency

Requires multiple donors or biological replicates measured under matched conditions. Applicable to the OPSCA, 300BCG, ParseBioscience, MSCIC, and SoundLife datasets.

#### SEM

Requires sufficient control samples to estimate GRN weights, and multiple unique perturbations to estimate TF shock vectors (*δ*). Applicable to all datasets except MSCIC, which lacks control samples, Nakatake, which contains only five controls, and 300BCG, where there is only one non-control perturbation condition (LPS).

#### Virtual cell

Requires control and multiple unique perturbation samples to predict perturbed states. Applicable to all datasets except the MSCIC and SoundLife datasets, which do not have multiple unique perturbations.

#### TF binding

Relies on ChIP-seq TF binding ground truth. Applicable to all datasets except Nakatake and MSCIC, which lacks ChIP-seq coverage. All other datasets use cell types (PBMC, K562, HEK293T, HCT116) with available ChIP-seq data.

#### Gene sets recovery

Uses perturbation signatures to compute enriched TFs from gene sets. Applicable to all datasets.

In addition, we conducted a quality control to determine which evaluation metrics were appropriate for each dataset, where for each dataset-metric combination, we applied three criteria: (i) Positive Control should perform higher than Negative Control, (ii) variability across methods, quantified as the coefficient of variation (CV = (max - min) / mean) should exceed a minimum threshold of 0.2, and (iii) the maximum observed score across all GRN models for the given dataset should exceed both Negative Control as well as a global minimum (Extended Data Table 2). The global minimum ensures that metrics are not considered applicable when all methods perform at a trivially low absolute level, even if one method nominally outperforms the Negative Control. The global minimums are set as follows: 0.005 for predictive metrics based on R^2^ (Regression, Virtual cell, SEM); 0.05 for metrics based on F1 score (TF binding, Gene set recovery); 0.5 for Wasserstein distance-based metrics (WS precision and recall); 0.3 for Replicate Consistency; and 2.0 (t-statistic) for TF recovery precision and recall.

Metrics were retained for a dataset only if they satisfied both criteria, ensuring that selected metrics demonstrated sufficient discriminatory power between methods and achieved meaningful performance levels. This analysis was performed across all available datasets and metrics, with results summarized to identify the optimal metric-dataset pairings for subsequent benchmarking analyzes.

### GRN inference

In this section, we describe the details of GRN inference using different methods. Each inference method was run according to the official documentation, and default parameters were applied unless otherwise specified. For each GRN, we identified the top regulatory edges based on the respective protocols and converted them into a standardized `TF-target-weight` format for downstream analysis. Given that different methods define regulatory interactions differently, we used the most relevant coefficient to represent the strength of the interaction as the `weight`. For multiomics methods producing a `TF-peak-target-weight` format, we aggregated regulatory weights from TFs to target genes; similarly, for CellOracle, scPRINT, scGPT, and Geneformer, which produce separate GRN models for individual cell populations in datasets based on primary cells (OPSCA, MSCIC, ParseBioscience, 300BCG, and SoundLife datasets), we averaged regulatory interactions across shared TF–gene pairs to generate a single GRN model per dataset (Supplementary Note 3). Also, for methods that do not natively filter for known TFs, we used curated TFs from [61] to filter the inferred GRNs.

#### scGLUE

We ran scGLUE [11] as described in the official documentation ^2^.

##### Prior regulatory graph and CRE-gene connections

We preprocessed both scRNA-seq and scATAC-seq data as instructed. A guidance graph was created, where peaks were associated with genes if they overlapped the gene’s body or had a distance of 150kb from the promotor region. GENCODE (V45) [82] was used to annotate genes which resulted in 17k annotations out of 22k genes. Genes without annotation were removed to comply with the rest of the pipeline. Subsequently, a GLUE model was trained to integrate the two types of omics data. Highly-variable genes (HVGs) were determined based on the recommended method and were used for the integration process. However, we used the entire genes to obtain the CRE-gene connections and final TF-gene connections instead of HVGs. The quality of the integration passed the defined threshold of 0.05 across all `n-meta` values. Genes and peaks were then embedded using the trained model, and cis-regulatory relationships between peaks and genes were obtained. To guide the inference process and reduce false positive rates, a skeleton was defined, limiting the search space based on the cosine similarity between peaks and genes in the embedded space. A threshold of 0.05 was defined for the adjusted *p*-values of the cosine score to draw significant regulatory connections.

##### TF-gene regulatory network

Initially, a preliminary TF-gene network was built based on co-expression using `pyscenic grn` [10] with the default settings (GRNBoost2). Next, the genome was scanned to connect the peaks with the motifs based on genomic overlap. We experimented with both of the given databases on JASPAR [83] and Encode [84] and decided on the latter as it slightly increased the size of the final network. By considering both gene-peak connections and peak-TF connections, TF-gene ranking was inferred using the `cis-regulatory-ranking` function. Additionally, we also included the proximal promoter regions around TSS in the generation of the regulatory feature database, as recommended in the instructions. This is to mitigate the issue of losing genes with no identified regulatory peak. Finally, `pyscenic ctx` was used to prune the preliminary network using the feature regulatory databases. We experimented with the parameters of this function to retain the largest network size as the default parameters yielded a very small network size (a few hundred).

##### Scenic+

We ran Scenic+ [12] as described in the official documentation ^3^.

##### Topic modeling

`Cistopic` was employed for topic modeling, which receives peaks as input and determines topics, defined as clusters of co-accessible regions. It resulted in an optimal number of 48 topics for the data at hand. The regions belonging to a topic are considered potential enhancers. `Cistopic` also uses co-accessibility scores between regions to handle dropouts in peak data.

##### TF motif enrichment analysis

`Cistarget` was used to detect enriched TF motifs per topic. For this, `Cistarget` uses a method named recovery curve to identify the enriched motifs for the sets of regions given in a topic, as well as to identify the top regions connected to each motif. We used a precomputed dataset suggested by Scenic+ for region-motif connections required in the `Cistarget` analysis.

##### TF-gene and region-gene scores

Region-gene scores were determined using gradient-boosting regression, predicting gene expression from region accessibility within a gene’s search space, which spans 1 kb to 150 kb around the gene or to the nearest gene’s promoter. The promoter region is defined as ±10 bp around the transcription start site. The obtained importance scores served as region-to-gene scores. TF-gene scores were derived similarly, predicting TF expression from gene expression counts, with gene importance scores indicating TF-gene relevance. Positive and negative interactions were distinguished using Spearman correlation for both TF-gene and region-gene pairs.

##### eRegulon creation

Region-gene connections were binarized using (0.7, 0.80, 0.90) as quantiles and (10, 15, 25) as the top number of regions per gene with a `rho-threshold` of 0. These parameters, chosen by consulting the authors of Scenic+, are less strict compared to the default settings, and retained more regulatory elements, whereas the default parameters resulted in a very small set of target genes and TFs. After, TF-region-gene triplets were formed by linking the remaining connections to the identified TFs from the motif enrichment analysis. Next, enrichment analyzes were conducted using the TF-to-gene importance score to determine the top genes as the targets of eRegulon. eRegulons with under ten targets were discarded.

#### FigR

We ran FigR [24] as described in the official documentation ^4^.

##### CRE-gene association testing

For each gene, all peaks located within a 10 kb distance from its TSS were identified. Subsequently, the Spearman correlation between peak counts and gene expression across all cells is computed. To assess the statistical significance of each CRE-gene correlation, a significance analysis is conducted by randomly choosing 100 background peaks to derive a *p*-value. After accounting for FDR, a threshold of 0.05 was used to identify significant CRE-gene connections. 180 Domain of Regulatory Chromatin (DORC) genes were identified by a threshold of 10 peaks per gene. In the original tutorial, only DORC genes are used during GRN inference. However, we opted for using all genes to retain sufficient regulatory elements in the GRN.

##### TF-gene associations

The DORC accessibility score was calculated by summing the raw counts of peaks connected to each target gene. The DORC scores and RNA counts were smoothed using the KNN algorithm with 48 neighbors, which are the topic numbers inferred in the Scenic+ workflow. Using smoothed DORC accessibility and RNA count matrices, we determined enriched genes for different TF binding motifs, as well as the correlations between the RNA counts of target genes with TF genes. Then, we applied the 0.05 threshold to retain significant connections.

#### CellOracle

We ran CellOracle [3] following the official documentation^5^.

##### Base GRN construction

We utilized `Cicero (version 1.3.9)` to calculate co-accessibility scores between pairs of scATAC-seq peaks. Peaks intersecting TSS within 1 kb were identified using annotations from BioMart (http://sep2019.archive.ensembl.org). By applying a threshold of 0.8 on co-accessibility scores, we filtered for a shortlist of CRE-gene connections. Transcription factor (TF) binding sites were then extracted from Gimmemotifs (version 0.17.1) and integrated with the CRE-gene connections.

##### GRN inference

The scRNA-seq data was filtered, normalized, and log-transformed according to the guidelines provided. Missing counts were imputed using a KNN method with 50 nearest neighbors. We ran the pipeline including all target genes and filtered out insignificant genes with a *p*-value threshold of 0.05.

#### GRaNIE

GRaNIE was originally designed for bulk multiomics data. However, specific preprocessing steps have been introduced in its official documentation^6^ to make it compatible with single-cell data.

##### Data preprocessing

We followed the default preprocessing steps outlined in the documentation using Seurat. For RNA, this involved a standard processing and normalization with `SCTransform` and dimensionality reduction using the first 50 principal components. For ATAC, a standard latent semantic indexing (LSI) normalization was performed (TF-IDF normalization, followed by `FindTopFeatures` using a cut-off of 0, and lastly a singular value decomposition was performed using `RunSVD`). The modalities were then integrated using the weighted nearest neighbor algorithm with `FindMultiModalNeighbors` with the PC components from RNA and LSI for ATAC. The integrated data was then clustered using Leiden with varying resolutions from 0.1 to 20. Clusters with fewer than 25 cells were removed, as recommended. Subsequently, the single-cell data was pseudobulked by calculating the mean count across cells of each cluster using the `AggregateExpression` function.

##### GRN inference

For each cluster resolution, we ran GRaNIE with the default settings unless noted otherwise. For the TF database, we used the INVIVO collection of HOCOMOCO v12 [85]. No further normalization was done as counts were already pre-normalized. Also, no additional filtering was performed for neither RNA nor ATAC. GRaNIE jointly infers TF-CRE and CRE-gene interactions based on the covariation of expression data and peak data. We employed Spearman’s correlation method, as recommended, to determine covariations. Two thresholds were applied to refine the inferred network: one at the TF-peak level (FDR 0.2) and one at the CRE-gene level (FDR 0.1).

#### GRNBoost2

GRNBoost2 method was applied using the `grnBoost2` function from the `arboreto (version 0.1.6)` Python package ^7^. The algorithm accepts a list of putative TFs as input, which we provided via the `tf names` parameter to reduce computation time.

#### Scenic

We ran `PyScenic` within the `aertslab/pyscenic:0.12.1 Docker` container and following the documentation ^8^. GRNs were inferred using the `pyscenic grn` command, with GRNBoost2 as underlying GRN inference algorithm, and using the same list of putative TFs. Inferred GRNs were next pruned with the `pyscenic ctx` command and using the `cisTarget` databases available at https://resources.aertslab.org/cistarget/databases. We modified the parameters to increase the retained number of edges by setting `rank threshold` to 10000 (5000 was the default).

#### PPCOR

Pairwise partial correlations were computed using the `pcor` function from the `PPCOR (version 1.1)` R package ^9^. We used the default correlation measure, namely the Pearson correlation coefficient (`method`=“`pearson`”). While `pcor` provides the correlation coefficient as well as the test statistic and *p*-value for each pair of genes, we relied on the correlation coefficients directly. Nonetheless, we suggest that correlation coefficients and test statistics can be used interchangeably since the Pearson test statistic is a monotonic function of Pearson correlation when the sample size is fixed.

#### Portia

`Portia (version 0.0.21)` was run as described in the README file at the root of the GitHub repository ^10^, except that the Box-Cox transform was disabled (`method`=“`no-transform`”) to alleviate potential numerical stability issues related to dropout. Similarly to GRNBoost2 and Scenic, the curative list of known regulators was provided (via the `tf idx` parameter).

#### scPRINT

scPRINT was run using the example notebook provided in its GitHub repository ^11^, in consultation with its authors. We experimented with various parameter settings, including different values for `num genes` (e.g., 5000 and 10,000), `max cells` (1000, 2000, and 5000), and different model checkpoints (*medium* and *large*). Since scPRINT requires single-cell count data as input, its evaluation was limited to the OPSCA, Norman, and Replogle datasets, as the other datasets were only available in normalized (pseudo)bulk format.

#### scGPT

We used the latest GitHub version (commit #5491783) together with the base model weights available on Google Drive (https://drive.google.com/uc?id=1MJaavaG0ZZkC_yPO4giGRnuCe3F1zt30), referred to as `best model.pt` (149M parameters). The gene–gene interaction network was derived from the mean attention matrices at layer 11 (the final layer) of the model. The set of genes, the number of cells per cell type, the aggregation procedure, and the conversion of attention values into a ranked list of gene–gene interactions were performed identically to those described for scPRINT.

#### Geneformer

We used the latest Hugging Face version (commit #91215c4) with the `Geneformer-V2` model (104M parameters) available in the same repository. The gene–gene interaction network was obtained by averaging attention matrices across all layers of the model, analogous to the approach used for scPRINT. In contrast to scGPT, for `Geneformer` only genes expressed in a given cell were considered; the attention matrix for a cell was set to zero for non-expressed genes. The gene set, cell type–specific cell counts, aggregation procedure, and transformation into a ranked list of top gene–gene interactions were performed as in scPRINT.

### Baseline control models

In addition to the inferred GRNs, we created three control models: a Negative Control, a Positive Control, and a baseline model. For the Positive Control, we computed pairwise Pearson correlations between all genes using both the inference and evaluation datasets. To reduce false positives, we limited the regulators to known TFs extracted from [12]. We then retained the top 50,000 regulatory edges based on their weights. Since the perturbation dataset was also used for GRN evaluation, we expected the Positive Control to exhibit high performance in the regression analysis. The baseline model was created similarly, with the exception that Pearson correlations were calculated using the inference dataset. For the Negative Control, we assigned binary regulatory weights randomly while maintaining a sparsity rate of 0.98, consistent with the inferred GRNs. The number of TFs was matched to the average TF count across the inferred GRNs, and target genes were sampled from the entire set of genes under evaluation.

### Overall scoring of GRN inference performance

To summarize GRN inference performance across datasets and metrics, we computed an overall score using a three-step aggregation procedure. First, scores were converted to ranks across GRN inference methods for each dataset–metric combination. Second, for each method, we took the median normalized rank across all applicable dataset–metric combinations, computed separately within multiomics (2 datasets, 11 dataset–metric pairs) and transcriptomics (7 datasets, 57 dataset–metric pairs) settings. Finally, the overall score was computed as the mean of the two modality-level medians, using only the modalities for which a method was evaluated. This procedure ensures that (i) each dataset–metric combination contributes equally to the overall score through ranking, (ii) differences in metric and dataset applicability across methods are appropriately accounted for, and (iii) multiomics GRN inference methods are not penalized for not being applicable to transcriptomics data.

### Performance analysis of context specific GRNs versus global models

To assess the performance of pre-existing global GRN models across different biological contexts, we compared their predictive accuracy against dataset-specific networks inferred de novo. Global GRN models, sourced from PEREGGRN [44], were evaluated alongside networks generated by standard inference methods applied to each individual dataset. For each evaluation dataset, all available global models were loaded and reformatted to match the dataset-specific gene space, ensuring fair comparison. Both global and dataset-specific networks were subjected to identical evaluation procedures using the full suite of applicable metrics for each dataset. Considering that the global models contain several cell types linked to PBMC, we analyzed context specificity across all our datasets linked to PBMC, namely OPSCA, 300BCG, ParseBioscience, and SoundLife datasets.

### Permutation analysis of GRN models

We conducted a series of perturbation analyzes on regulatory information in GRN models. Controlled perturbations were systematically introduced to TF-gene properties obtained from GRNs at varying intensities to assess their effects on performance metrics. For TF-gene connections, we shuffled the TF-gene edges at varying levels: 0%, 10%, 20%, 50%, and 100%. A 0% shuffle implies no change to the matrix, while 100% represents the complete permutation of the TF-gene edges. For the regulatory weights, we added Gaussian noise to the regulatory weights at varying intensities and evaluated the decline in the regression scores. The noise was sampled from a normal distribution 𝒩 (0, `intensity` × *σ*), where *σ* represents the standard deviation of the weights. For the regulatory sign, we randomly flipped the sign of regulation (e.g. positive to negative). For each perturbation level, we recalculated performance scores and evaluated the impact of the perturbations on performance decline. We run the experiments on a selected set of datasets (OPSCA, Replogle, ParseBioscience, and Norman) and GRN models (Scenic+, Scenic, Pearson Correlation, PPCOR, and scPRINT) to save computations.

### Causal evaluation of GRN inference

We conducted two experiments to assess whether our evaluation metrics prioritize causal regulatory relationships over purely correlational ones. In the first experiment, we flipped the direction of TF-gene interactions to analyze whether the performance is sensitive to the direction of regulations inferred by GRN models. We run this experiments for seven datasets (OPSCA, Replogle, Norman, MSCIC, ParseBioscience, and Xaira datasets) and all GRN methods. For each scenario, we calculated the performance score and compared it to the original GRN model. In the second experiment, we examined whether the Regression metric was sensitive to the type of regulators used to predict gene expression signatures. Specifically, we evaluated GRN model performance under two conditions: with and without filtering for known TFs. In the filtered setting, we restricted regulatory edges to those involving one of 1,800 curated TFs [12]. Given that the dataset contains approximately 20,000 genes, this filtering step reduces potential false-positive causal interactions by roughly 90%. We performed this experiment on the three largest perturbation datasets from Replogle and two datasets from Xaira, using GRNs inferred with the Pearson Correlation method. By comparing model performance across the filtered and unfiltered scenarios, we tested whether this metric preferentially reward regulatory relationships that are more likely to be causal rather than spurious correlations.

### Analysis of the effect of data normalization on performance

To evaluate the effect of gene expression normalization strategies on GRN inference performance, we conducted a systematic comparison across two normalization approaches. For each dataset, GRNs were inferred using different methods applied to two normalization schemes: (i) shifted logarithmic approach (SLA), representing the default preprocessing method in our pipeline, and (ii) Pearson residual (PR) normalization, which accounts for technical variation while preserving biological signal. For both approaches, we used `Scanpy` [86]. All inference methods were executed with identical parameters across normalization conditions, with the only variation being the input data layer used for network construction and GRN evaluation. The resulting GRN predictions were evaluated using the complete set of applicable metrics for each dataset.

### Analysis of GRN models structural properties and gene-, TF-wise performance

To analyze whether there is a common sets of regulatory interactions leading to higher performance, we compared topological and predictive features across four models (GRNBoost2, Scenic+, Pearson Correlation, and PPCOR) on the OPSCA dataset. Jaccard similarity was calculated for inferred TF–gene edges, as well as for the sets of inferred TFs and target genes.

To investigate GRN inference performance in a more granular way, gene-wise and TF-wise performance scores were derived from the Regression and WS distance metrics, respectively. Gene-wise scores were obtained from OPSCA dataset and TF-wise from Replogle dataset. Genes and TFs were stratified into easy-to-score (top 0.8 quantile) and hard-to-score (bottom 0.2 quantile) groups based on mean scores across models. For each gene, mean expression and expression variability (standard deviation) were computed from log-normalized pseudobulk profiles across all samples and related to per-gene mean *R*^2^ to assess whether expression level or variability drives model predictability. Gene set enrichment analysis was performed separately for easy and hard gene sets using `Enrichr` (via `gseapy`) querying `GO Biological Process 2023`, `KEGG 2021 Human`, and `Reactome 2024 databases`. Results from all three databases were pooled and filtered at an adjusted p-value threshold of 0.05. For TFs, mean WS distance across all GRN models was used to assign easy- and hard-to-score groups using the same quartile cutoffs. The global perturbational effect of each TF was summarized as the absolute log_2_ fold-change of genome-wide expression upon knockdown compared to control and related to mean WS distance to evaluate whether perturbation strength explains TF predictability.

### Analysis of regulatory relationship stability across perturbations and donors

To evaluate the stability of GRN inference, we analyzed TF-gene pairs inferred by four models—GRNBoost2, Scenic+, Pearson Correlation, and PPCOR—on the OPSCA dataset and re-estimated their regulatory weights across different sample selections. Specifically, we performed a five-fold sampling of the perturbation dataset; for each fold, regulatory weights were estimated using ridge regression, fitting one model per gene with the inferred TFs as predictors, and coefficients were standardized by the number of regulators per gene using the consensus set from the Regression metric. As a baseline, random GRNs preserving TF degree distributions were generated. Stability across perturbation samples was quantified as

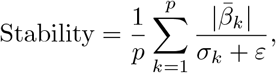

where 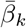 and *σ*_*k*_ are the mean and standard deviation of TF *k*’s coefficient across folds, and *ε* prevents division by zero. Higher stability indicates that TFs consistently receive strong, reproducible weights. Stability distributions were compared between inferred and random GRNs, and across models, using the `Mann{Whitney U test`. A similar analysis was performed using donor-based splits, selecting two donors out of three to assess reproducibility across biological replicates.

### Evaluating the effect of imputation and pseudobulking on performance

Pseudobulking and imputation are commonly employed in single-cell analysis and GRN inference pipelines to mitigate data sparsity and improve signal-to-noise ratios. Here, we systematically evaluated the impact of these preprocessing steps on GRN inference performance using our evaluation metrics. To assess the effect of data aggregation, we conducted two complementary experiments. First, we performed meta-cell aggregation by clustering single-cell data using Leiden clustering at resolutions ranging from 1 to 19 [5]. For each resolution, we generated aggregated expression profiles and evaluated GRN performance alongside the original single-cell data. This analysis was performed on three single-cell datasets (OPSCA, 300BCG, SoundLife:Vaccine, and Norman). GRNs were inferred using Pearson Correlation and GRNBoost2. Due to computational constraints, GRNBoost2 was applied to a subset of resolutions (single-cell, 9, and 19).

Second, we evaluated per-group pseudobulking by comparing GRNs inferred from single-cell expression data to those derived from pseudobulk profiles. Pseudobulk profiles were generated by aggregating cells based on available biological annotations, including cell type, donor, well, and perturbation condition. This analysis was conducted on three large genetic perturbation datasets (Xaira:HEK293T, Xaira:HCT116, and Replogle). For each dataset, we constructed two input representations: (i) single-cell expression matrices preserving cellular heterogeneity, and (ii) pseudobulk expression profiles aggregated within each perturbation condition. GRNs were inferred using Pearson Correlation with identical parameters and transcription factor sets across both representations.

To evaluate the impact of imputation, we applied two methods: KNN imputation (implemented in *scikitlearn*) and `MAGIC` [87]. This analysis was performed on the same five single-cell datasets used in the metacell experiment (OPSCA, 300BCG, SoundLife:Vaccine, and Norman). GRNs were inferred using Pearson Correlation, and performance was compared between imputed and original single-cell data.

### GRN inference evaluation in the context of multiple myeloma precursor disease

We obtained a paired single-cell multiomics dataset (scRNA-seq and scATAC-seq) of primary bone marrow from 12 patients with myeloma precursor disorders (MGUS/SMM) enrolled in the NUTRIVENTION clinical trial, deposited under GEO accession GSE311602 [88]. The dataset comprises cells profiled using two protocols: 10x Genomics Multiome and DOGMA-seq. The raw data consisted of approximately 101 000 cells, 36 000 genes, and ∼1.3 million peaks. Peaks were merged using `bedtools` [89] to generate 271 470 consensus peaks. Quality control retained cells with at least 200 detected genes and between 500 and 100 000 peaks, and genes detected in fewer than 10 cells were removed. Expression data were normalized using a shifted logarithmic transformation, consistent with other datasets. The data were then split by protocol, yielding a Multiome subset (12 796 cells × 21 826 genes, 3 donors) and a DOGMA-seq subset (25 000 cells × 21 826 genes; top 3 donors selected to retain 25 000 cells). GRN models were inferred using five methods—GRNBoost2, PPCOR, CellOracle, Scenic+, and GRaNIE—resulting in ten GRNs in total; for each model, the top 50 000 regulatory edges were retained, consistent with our evaluation pipeline.

We defined a set of 31 canonical TFs defining immune cell identity (`Immune core TFs`), curated from the literature, spanning major immune lineages present in the bone marrow microenvironment: myeloid and dendritic cells (SPI1, IRF8, CEBPA/B, MAFB); B cells and plasma cells (PAX5, EBF1, BACH2, BCL6; PRDM1, XBP1, IRF4); T cell subsets (TBX21, GATA3, RORC, FOXP3, BATF, TCF7, LEF1, BCL11B, ZEB2, ETS1, KLF2); CD8^+^/NK (EOMES, RUNX3, ZNF683); and pan-lymphoid progenitors (IKZF1, IKZF2, RUNX1, IRF7) [90, 41, 43, 42, 35, 33, 37]. We confirmed that all 31 TFs are expressed within the immune compartment, as supported by their inclusion in the CellTypist model [91].

For MMP-associated genes, we used the NHGRI-EBI GWAS Catalog [92] and filtered for the keywords *multiple myeloma, plasma cell myeloma, myeloma, MGUS, monoclonal gammopathy*, and *smoldering myeloma*, retaining only genome-wide significant associations (*p <* 5 × 10^−8^), yielding 128 genes. For MMP-associated TFs, we combined GWAS results with manually curated TFs from the literature. For GWAS, we further filtered the 128 genes for genic SNPs (those falling within a gene body, excluding intergenic and synonymous variants), intersected with a curated list of 1,638 human TFs [61], and retained only TFs annotated to immune-related Gene Ontology biological processes (316 GO terms, including immune response, lymphocyte activation, leukocyte differentiation, and related categories; MSigDB C5 [76]), resulting in 4 TFs:

*KLF2, PRDM1, SP140*, and *SP3*. We curated an additional 15 TFs from published MM biology, spanning the MYC pathway (*MYC, MYCN, MAX*), NF-*κ*B family (*NFKB1, NFKB2, RELA, RELB*), Ikaros family (*IKZF1, IKZF3*), MAF family (*MAF, MAFB*), and key oncogenic regulators (*IRF4, STAT3, HIF1A, TP53*) [38, 93, 39, 33], yielding a combined set of 19 `MMP TFs`.

To analyse whether annotated TFs are represented in the network, we calculated out-degree centrality for each GRN model and dataset and assessed the overlap of top 20 central TFs with the annotated TFs. For the representation analysis of annotated genes, we calculated in-degree centrality per network and dataset and compared the centrality of annotated genes to other genes in the network, where we used a Mann–Whitney U test to assess statistical significance.

## Supplementary Figures

**Supplementary Figure 1:**
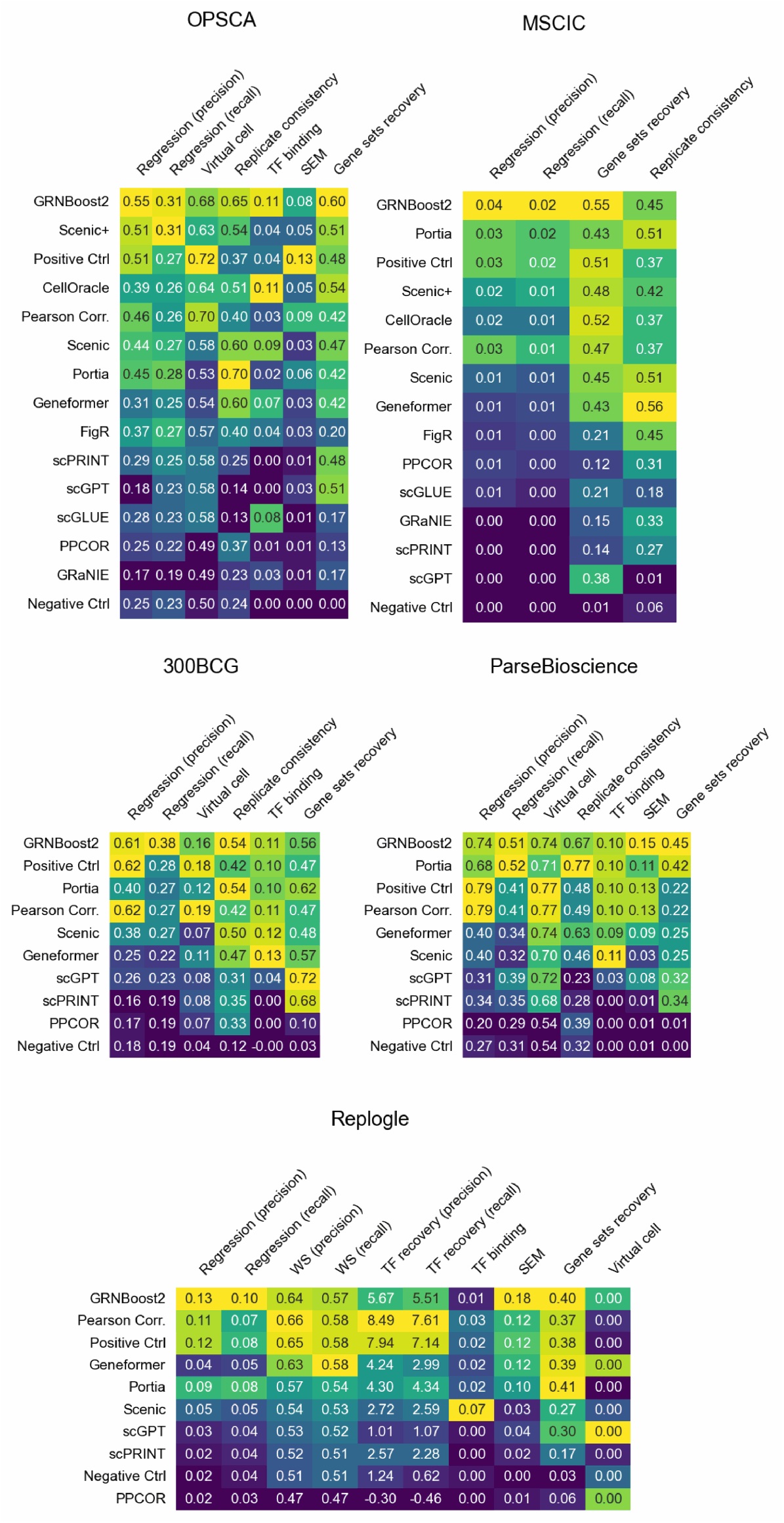
Performance scores

**Supplementary Figure 2:**
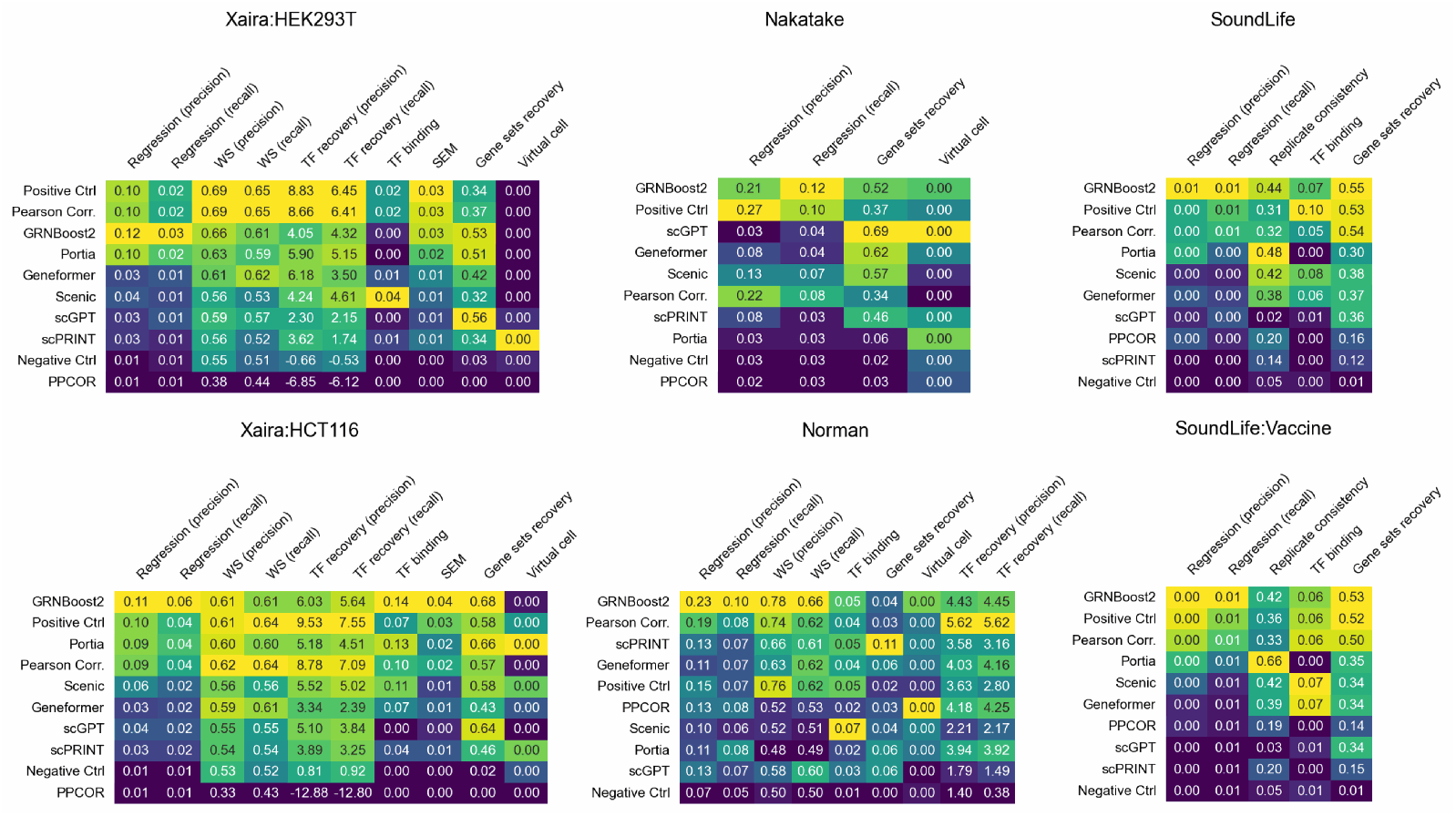
Performance scores

**Supplementary Figure 3:**
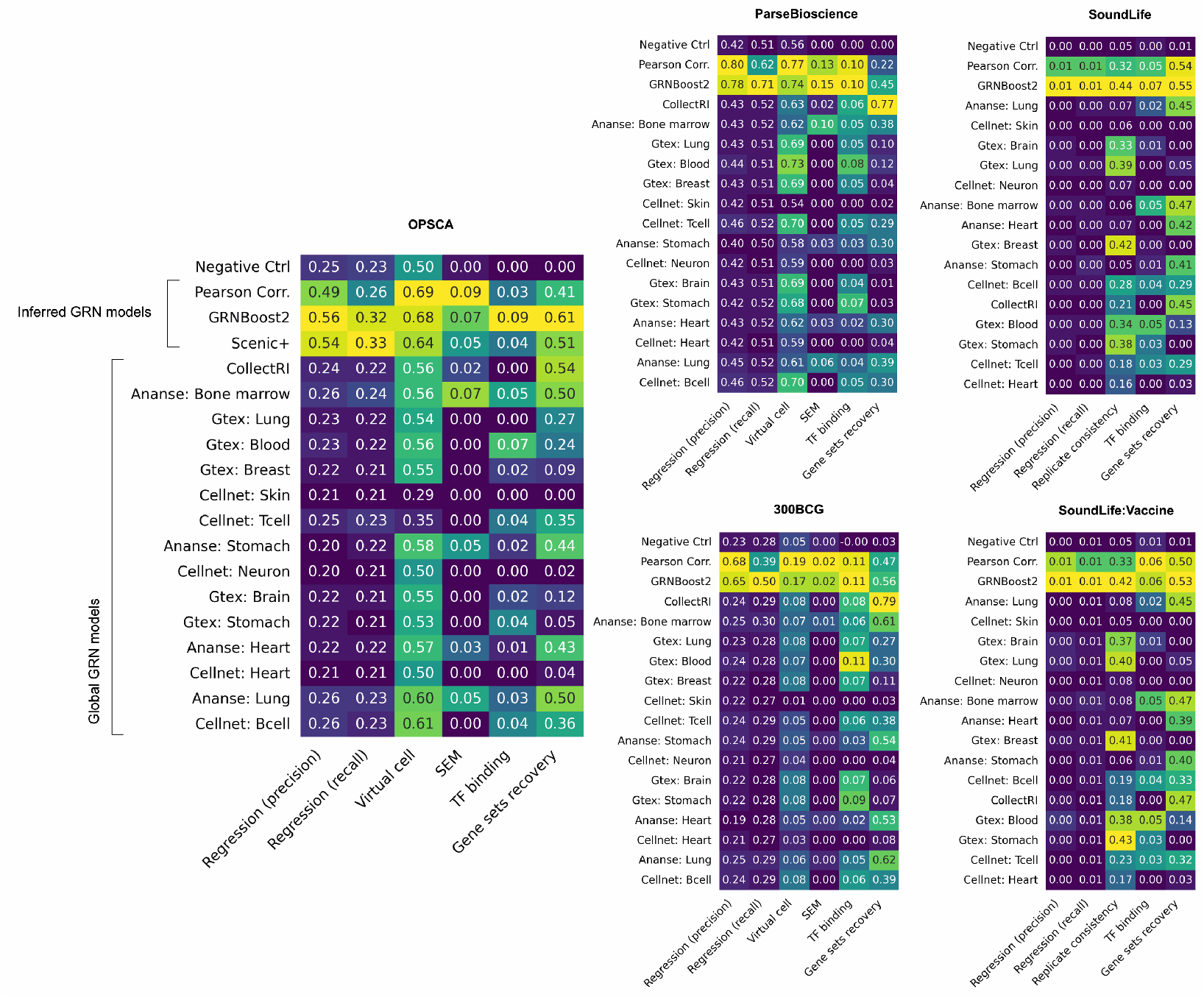
Performance comparison of inferred GRN models versus global (non-context specific) models.

**Supplementary Figure 4:**
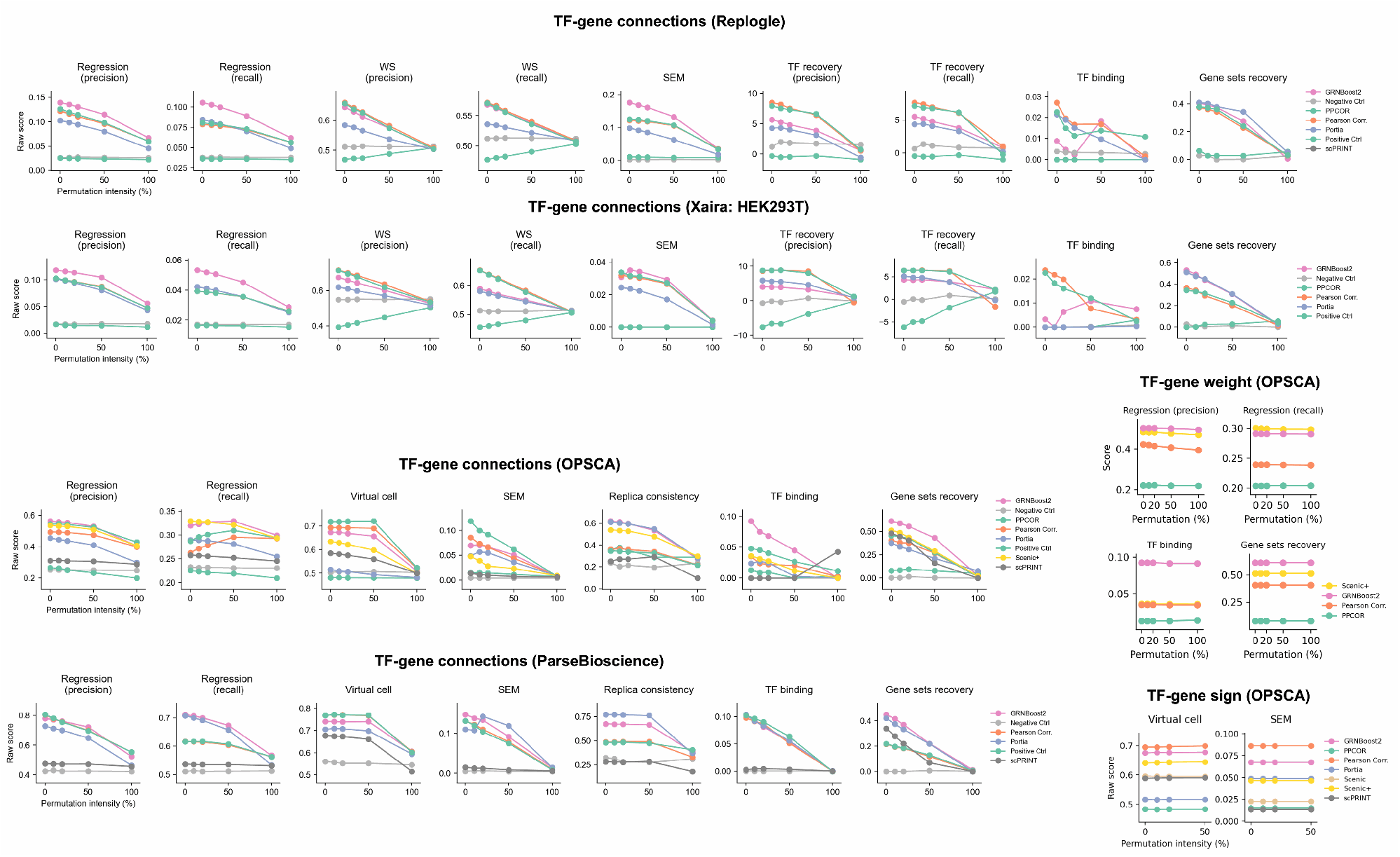
Performance analysis of the permutation of GRN connections, weight, and sign. While TF→gene connections showed notable drop across all metrics and datasets, the weight and sign permutation was either not applicable for certain metrics or did not yield any notable drop. We only provided one dataset as example.

**Supplementary Figure 5:**
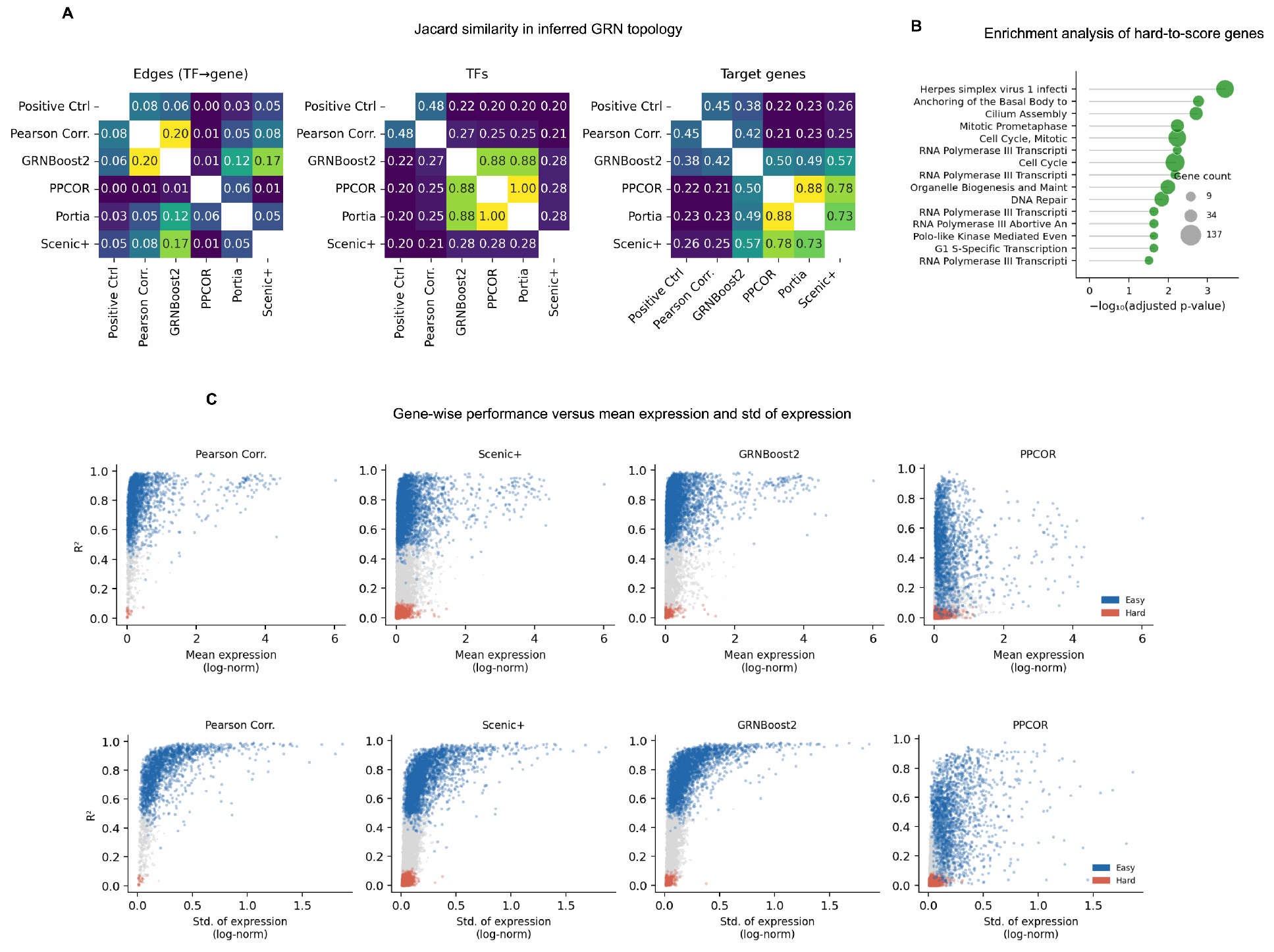
(A) Jaccard similarity calculated on different topological features of GRN models. (B) Results of enrichment analysis for hard-to-score genes in the Regression metric. (C) Performance of TF-wise scores obtained from WS distance metric versus the perturbation strength of each TF, calculated as the log gold change against control samples.

**Supplementary Figure 6:**
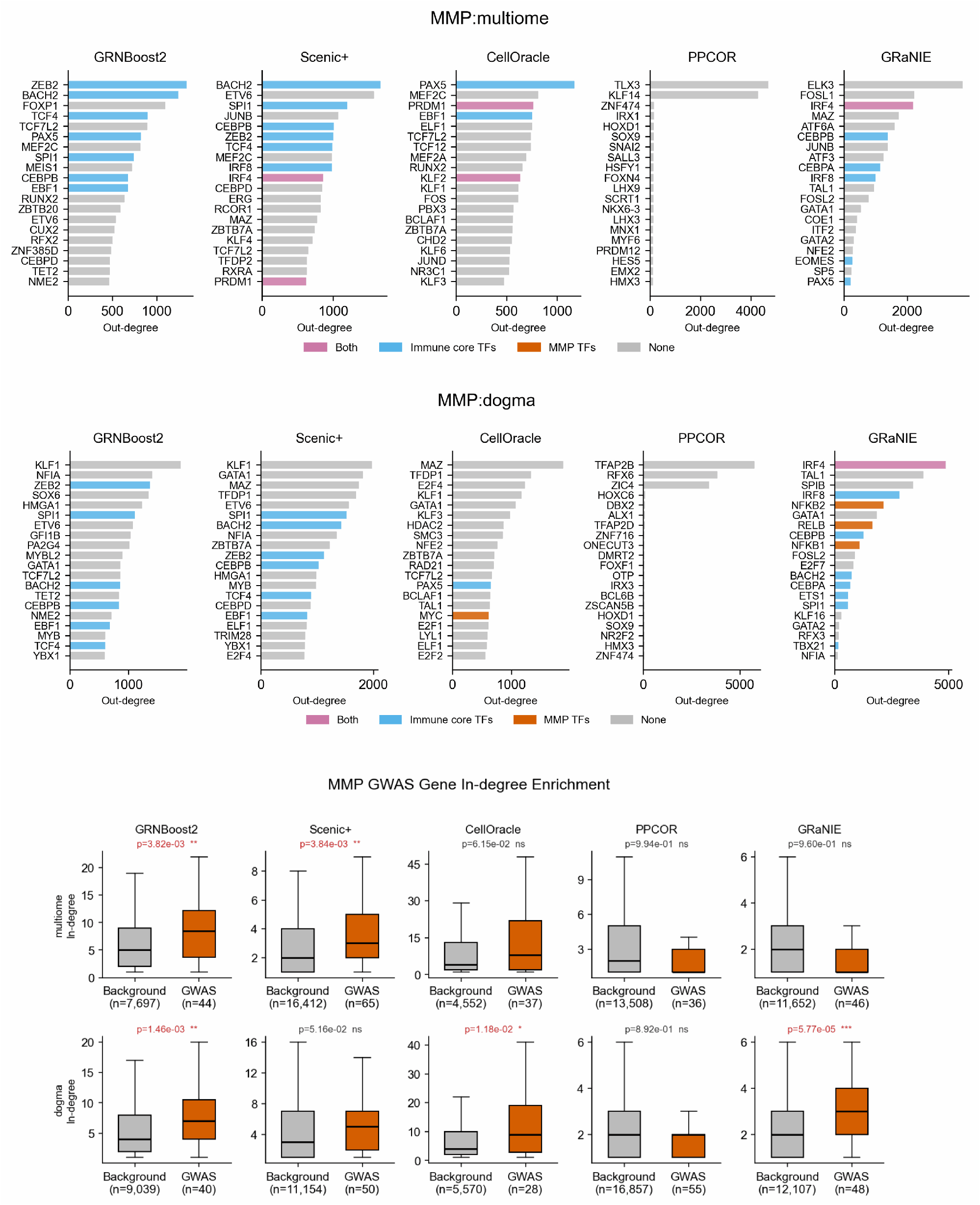
(Bar plots) Out-degree centrality (top 20) obtained for different GRN models across two sequencing technology of 10x Genomics Multiome and DOGMA. Those overlapping with core immune markers and MMP-associated TFs are annotated. (Box plots) Results of the rank test to determine whether MMP associated genes were more central (in-degree) than other genes in the network.

**Supplementary Figure 7:**
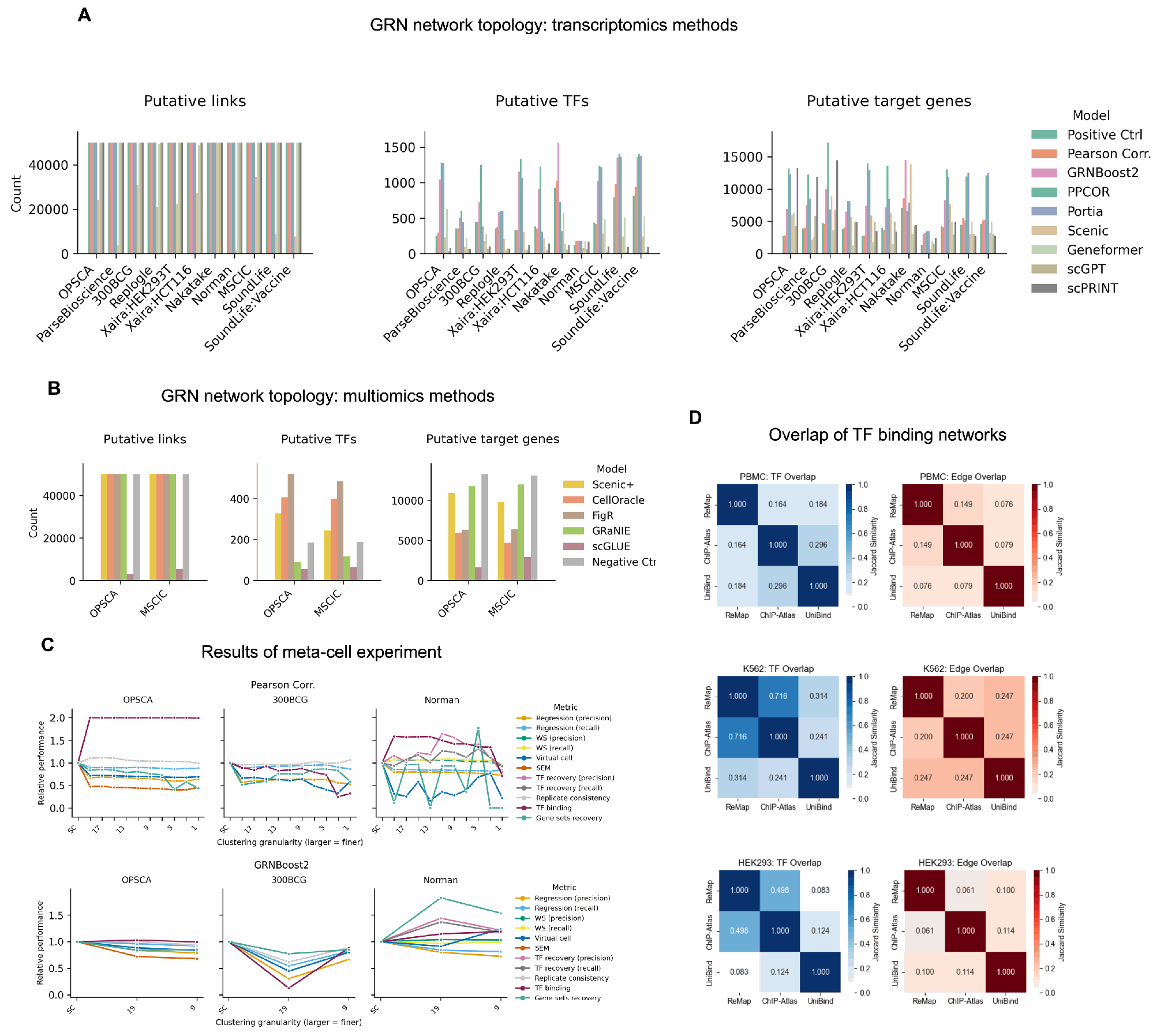
(A and B) Topological statistics of GRN models for different datasets, transcriptomics methods (top) and multiomics methods (bottom). (C) The effect of granular data aggregation on GRN inference performance for two methods of Pearson Correlation and GRNBoost2. Original single cell (SC) data are the most left data points, while the following numbers indicate the Leiden resolution used to aggregated the single cell data, with bigger the number, higher the resolution. (D) The overlap of TFs (left panels) and edges (right panels) between ground truths constructed from different TF binding data sources across different cell types.

**Supplementary Figure 8:**
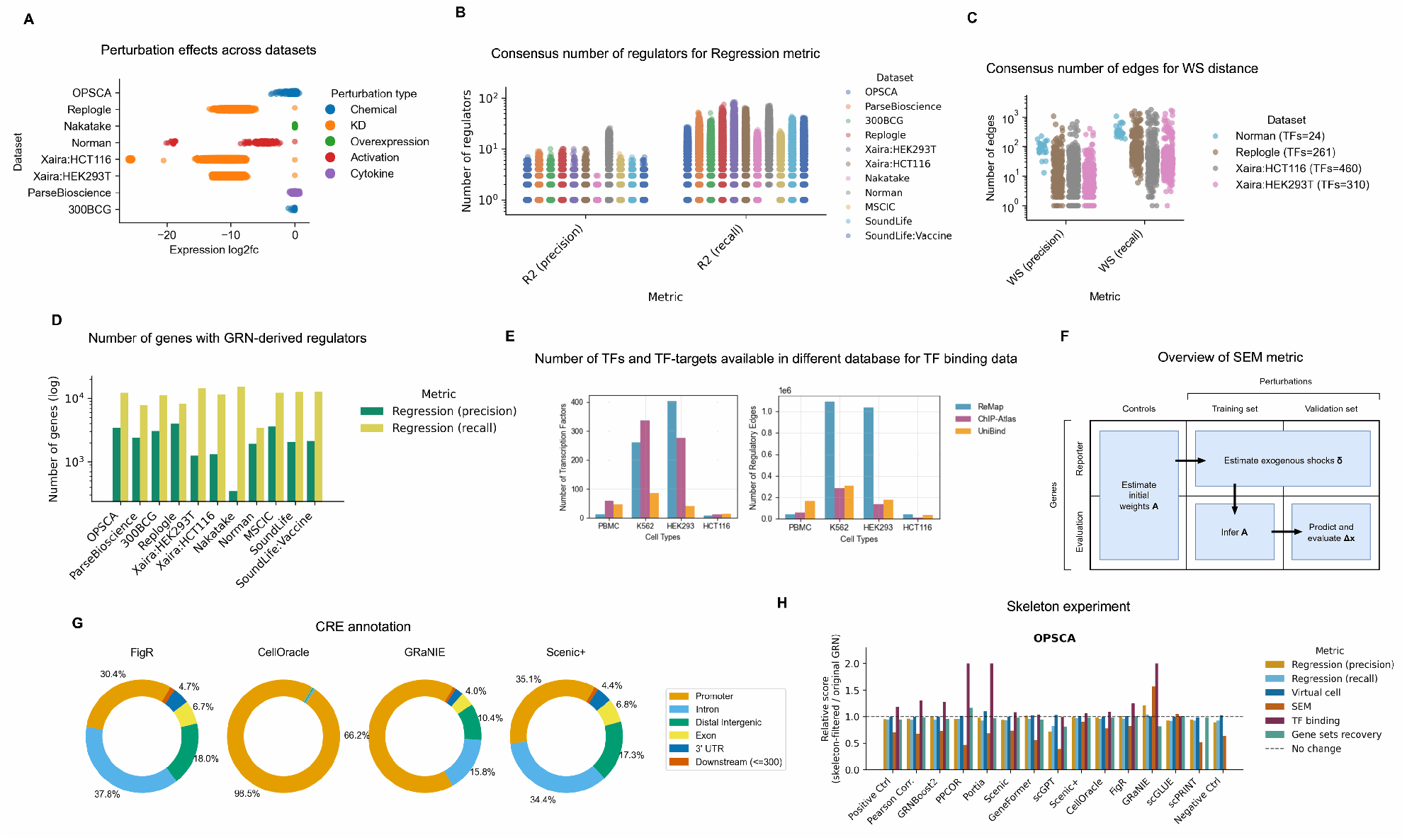
**(A)** The perturbation effect calculated as the signed log fold change of each perturbed sample versus control. **(B)** Distribution of the number of regulators for different metrics of regression. **(C)** Distribution of the number of edges for WS distance. Each dot represents one TF. **(D)** The number of genes with GRN-derived regulators across different datasets for regression. **(E)** Statistics of ground truth obtained from Chip-seq data from different sources. **(F)** SEM-based GRN evaluation using perturbation data. The data is split into four sets to prevent information leakage and ensure that SEM estimates perturbations Δ*x* from unknown shocks *δ* without bypassing the network topology *A*. **(G)** Annotation of CREs resulted from different GRN models for OPSCA dataset. **(H)** Results of Skeleton filtering.

**Supplementary Figure 9:**
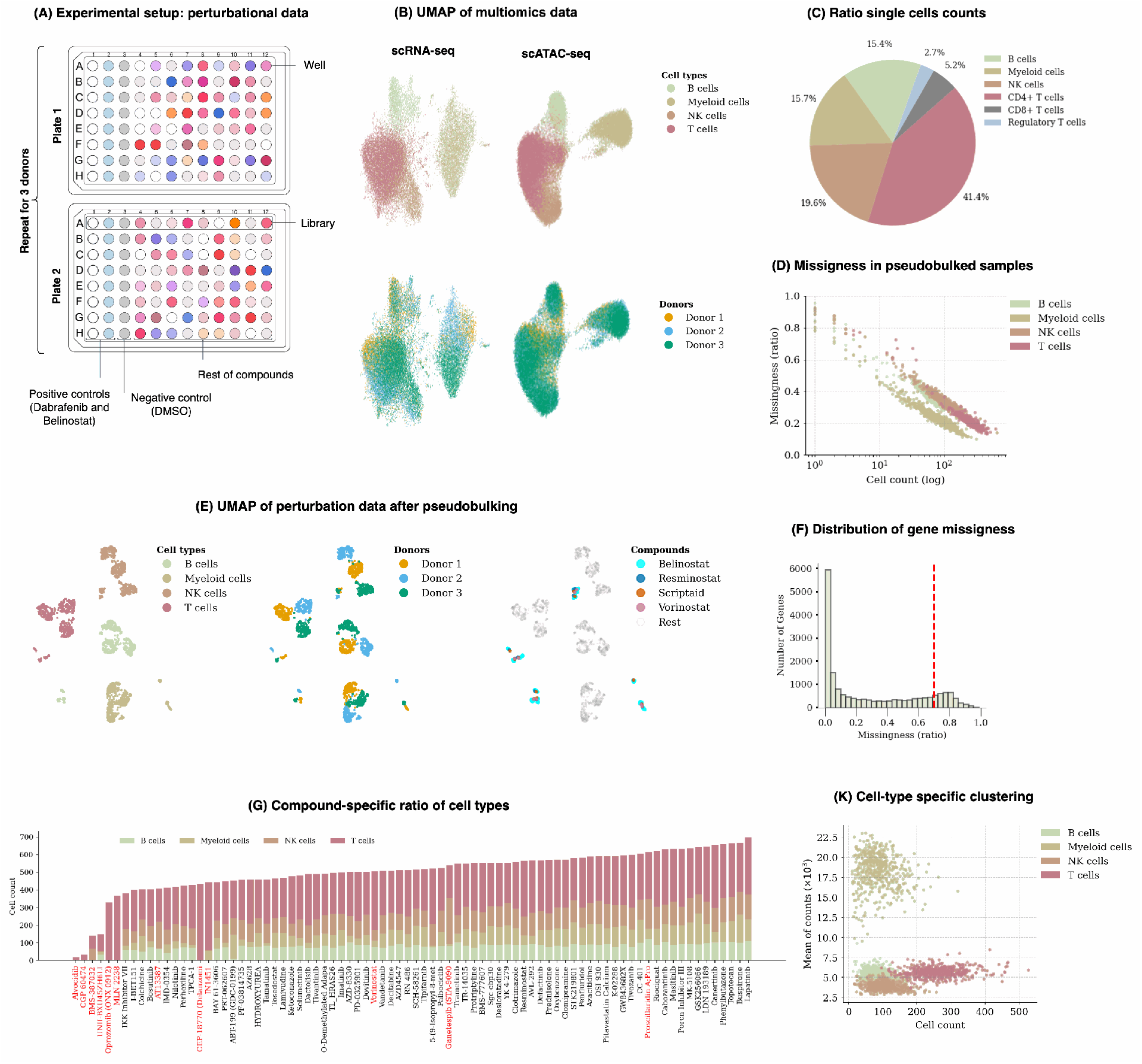
Preprocessing of OPSCA dataset. **(A)** Experimental design of perturbation dataset. **(B)** UMAP of the multiomics dataset used for GRN inference. **(C)** Cell type composition of the perturbation dataset. **(D)** Missingness of samples versus cell count, stratified by cell type. Missingness is calculated by summing the number of missing genes across the cells found in each pseudobulked sample. **(E)** UMAP of perturbation dataset after pseudobulking. **(F)** Distribution of gene missingness ratio across pseudobulked samples. The red line shows the cut-off threshold of 0.7 to filter low coverage genes. **(G)** Compound-specific composition of the perturbation dataset. Compounds in red have been identified as outliers in our preprocessing. Among the compounds that are neither controls nor outliers, only half are shown due to space restrictions. **(H)** Cell type composition of compounds that exhibited abnormal distribution on certain donors. **(K)** Sum of counts versus cell count of data pseudobulked by mean, stratified by cell types.

**Supplementary Figure 10:**
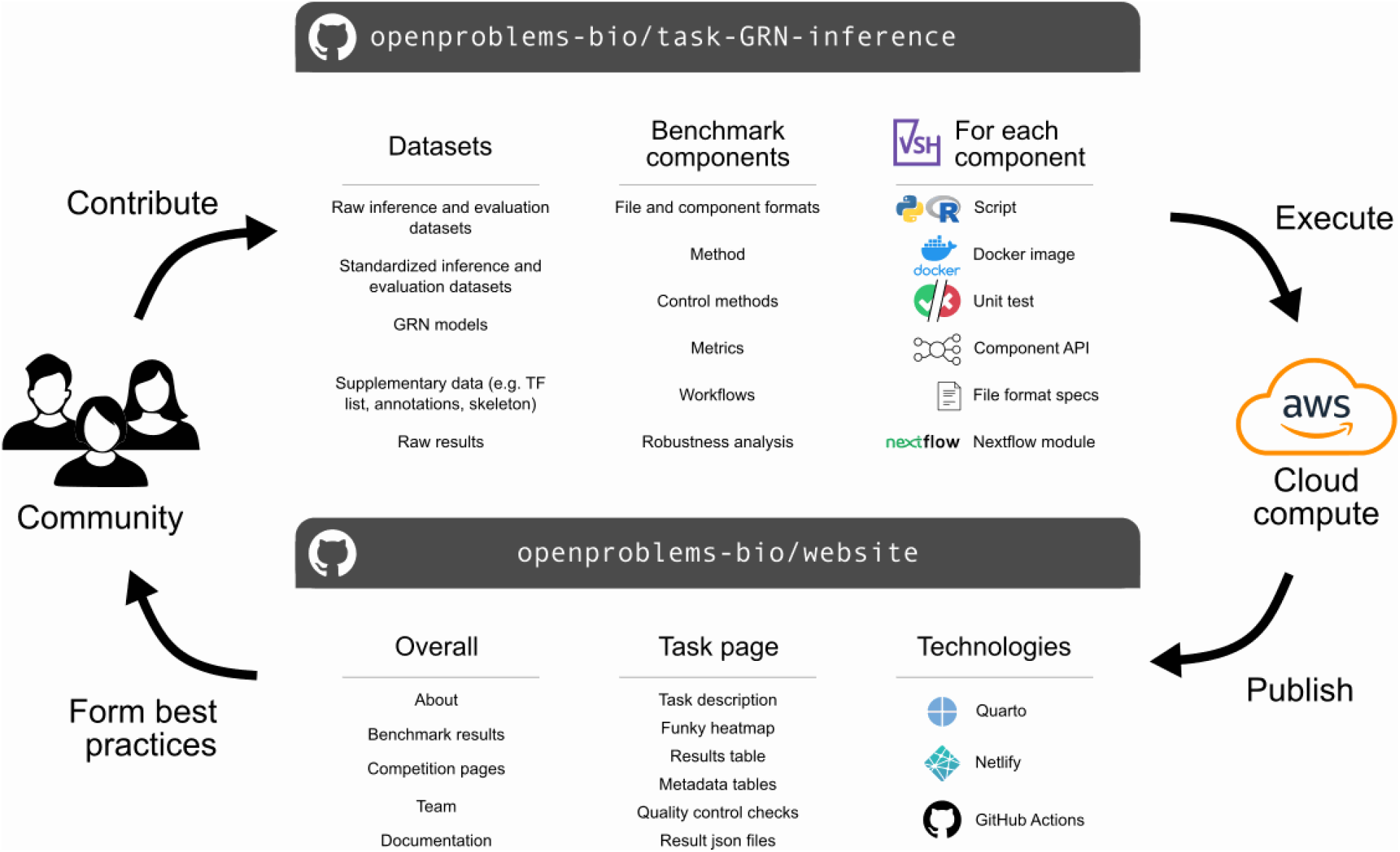
An overview of the *Open Problems* ecosystem where geneRNIB is hosted.

## Supplementary Note 1

geneRNIB provides a flexible and reproducible platform for integrating new GRN methods, datasets, and evaluation metrics. The platform leverages modern frameworks such as `Docker`, `Viash`, and cloud infrastructure to ensure reproducibility. Automated unit testing via `GitHub Actions` validates all components against test datasets before deploying, guaranteeing continuous integration and reliability. Inference and evaluation datasets are stored in the cloud, enabling consistent access and facilitating reproducible computations. Cloud-based execution also provides real-time monitoring of computational requirements for each GRN inference method, offering valuable insights into performance trade-offs. Benchmarking is repeated after every new integration to maintain up-to-date performance evaluations, and results are displayed on a dynamic leaderboard reflecting the latest developments. geneRNIB includes extensive documentation to streamline user interactions and foster accessibility. As part of *Open Problems*, the platform promotes an active support ecosystem to assist users with development and troubleshooting, ensuring a collaborative and inclusive experience for contributors.

The overview of geneRNIB’s components and pipelines is summarized in Supplementary Fig. 10. All standalone code—including GRN inference methods, evaluation metrics, preprocessing scripts, robustness analyzes, and workflows—is implemented as `Viash` components. Each component comprises a script file accompanied by metadata that defines its inputs, outputs, and software requirements, facilitating the creation of Docker images. To address the complexity of software dependencies for certain GRN inference methods, we generated individual Docker images for each method and published them on Docker Hub.

To ensure interoperability across components, the repository primarily uses the `AnnData` package [94] to establish the standard format for both input and output files. Strict formatting standards are enforced for all file types to maintain consistency. geneRNIB contains a major `Nextflow` workflow that runs all GRN inference methods and evaluation metrics and provides final scores. The workflows are executed on `AWS Batch`, enabling components to run in parallel according to workflow requirements. Different AWS instance types are employed based on the specific memory, CPU, and GPU requirements of each component, ensuring efficient resource allocation.

### Agentic geneRNBI: autonomous GRN inference, evaluation, and troubleshooting

geneRNIB includes a structured agent instruction file (AGENT.md) designed to enable AI coding agents to autonomously interact with and extend the benchmark. The file provides a comprehensive operational overview of the framework, including the three-stage pipeline architecture (inference → evaluation → postprocessing), execution environment decision guides for Docker/Viash, Conda, Singularity, and SLURM, task-specific routing to detailed documentation, standardized data contracts for all pipeline file formats, and a systematic troubleshooting protocol. We validated the effectiveness of this design by running the complete GRN inference and evaluation workflow end-to-end using an AI agent, as well as integrating new GRN inference methods into the benchmark through agent-assisted development. These experiments demonstrate that the provided instructions are sufficiently detailed for AI agents to contribute meaningfully to the benchmark with minimal human intervention.

## Supplementary Note 2

The experimental design of the perturbation dataset is depicted in Supplementary Fig. 9-A. PBMCs were arranged on 96-well plates, with two columns dedicated to Positive Controls (Dabrafenib and Belinostat) and one column to the baseline compound, dimethyl sulfoxide (DMSO), which served both as Negative Controls. The Positive Controls were selected for their known significant impact on transcription. The remaining wells on each plate were allocated to all the other 144 compounds, one per well. Each experiment was conducted with three donors, resulting in a total of six plates (two plates per donor). All samples within each row were pooled and sequenced. The Cell Ranger pipeline was used by the OPSCA competition organizers to process the reads [95]. In addition, PBMCs were cultured in T75 flasks and treated with DMSO, followed by the generation of integrated scRNA-seq and scATAC-seq data using the 10x Multiome assay. Computational methods were employed by the same organizers to annotate cell types in both the multiomics and perturbation datasets.

The UMAP visualization of the perturbation and multiomic datasets are given in Supplementary Fig. 9-B with a clear clustering of the cell types for both data types. The single-cell perturbation data and multiomics data were originally preprocessed by the OPSCA competition organizers to remove low-quality cells, genes, and peaks. We used the multiomics data without further preprocessing as the inference data. However, certain biases remained in the perturbation dataset, as highlighted in our report [96] and in reports from other contenders [97]. Therefore, we conducted careful preprocessing to address these identified biases.

The original perturbation data exhibited a high imbalance among different cell types in terms of cell counts, as shown in Supplementary Fig. 9-C. CD4+ T cells constituted approximately 41% of all cells, whereas T regulatory cells comprised only 2.7%. This imbalance has been shown to result in heteroscedastic behavior, exhibiting cell type-specific variations in the resulting training data [96]. To address this, we merged all three types of T cells into a single cell type in both perturbation data as well as multiomics data, which improved the balance in cell type ratios. Next, we eliminated compound-cell type pairs with fewer than 10 single cells, as they resulted in significant missingness (Supplementary Fig. 9-D) and were associated with abnormal observations in previous reports [96]. Next, we removed the genes that were absent in more than 70% of samples (Supplementary Fig. 9-F), to assure that the included genes are sufficiently represented among the samples. This filtering reduced the gene space to 15,215. As a result of this filtering and those mentioned earlier, the missingness (ratio of zero counts) in the pseudobulked dataset improved from the original 42% to 18%. The UMAP of pseudobulked samples are shown in Supplementary Fig. 9-E, with four compounds of Belinostat, Resminostat, Scriptaid, and Vorinostat showing strong effects in expression signatures.

In addition, we identified outlier compounds that either (i) exhibited excessive toxicity leaving only very few cells alive or (ii) caused significant shifts in cell type ratios. Supplementary Fig. 9-G illustrates the distribution of cell type ratios across different compounds, averaged across the three donors. Notably, four compounds-Alvocidib, UNII-BXU45ZH6LI, CGP 60474, and BMS-387032-demonstrated highly toxic effects, resulting in a significant reduction in cell counts. Consequently, we removed these compounds from the dataset. Additionally, we excluded four compounds—CEP-18770 (Delanzomib), IN1451, MLN 2238, and Oprozomib (ONX 0912) due to their substantial distortion of cell type ratios. Furthermore, we removed AT13387 and Ganetespib (STA-9090) for donor 3 only, and Vorinostat for donor 2 only, as they exhibited normal ratios in the remaining donors.

Overall, we observed a high expression signature for Myeloid cells compared to other cell types (Fig. 9-K).

## Supplementary Note 3

### Guidance on GRN model construction

The geneRNIB evaluation pipeline accepts GRN models in a standardized format consisting of TF, target gene, and edge weight. While transcriptomics-based GRN inference methods typically output TF–gene pairs, multiomics approaches often produce extended regulatory structures, such as TF–CRE–target-weight links. To harmonize these representations, we reformatted multiomics GRNs by retaining only TF–target gene connections, aggregating edge weights across intermediate regulatory elements. It is important to note that edge weights produced by different GRN inference methods may encode distinct biological meanings, including regulatory effect sizes, confidence scores, or attention weights in foundation models.

In the geneRNIB evaluation framework, these weights are interpreted primarily as relative confidence measures or the likelihood of a true regulatory interaction. Accordingly, edge weights influence multiple stages of our evaluations: the selection of the top 50k edges to standardize evaluation, which defines network connectivity for SEM and Virtual cell metrics, the identification of top regulators per gene for the Regression metric, and weighting TF–gene associations when computing TF activity scores for TF recovery. Therefore, users should carefully consider the interpretation and scaling of edge weights when constructing GRN models for evaluation with geneRNIB, ensuring that higher weights consistently reflect stronger or more confident regulatory relationships across GRN model.

Currently, our evaluation framework expects one GRN model per dataset. The rationale for this design is twofold. First, certain GRN inference methods, such as Scenic+, leverage variability across cell subpopulations during network inference, thereby producing a single GRN model for the entire dataset. Evaluating cell type–specific GRNs together with dataset-level GRNs would require methodological inconsistencies, including differences in cross-validation design, leading to results that are not directly comparable. To maintain standardized evaluation across methods, we therefore constrain the analysis to one GRN model per dataset. Second, several evaluation metrics, including Regression, SEM, and Virtual Cell, require sufficient sample sizes for robust model training and prediction. Splitting datasets into smaller cell subpopulations would substantially reduce statistical power and potentially diminish the robustness of these machine learning–based evaluations. Consequently, our framework requires a single GRN model per dataset. When multiple GRN models are provided, we aggregate them by averaging regulatory weights across shared TF–gene pairs.

### Potential limitations of precision matrices in GRN inference

Among single-modality inference methods, our analysis shows that PPCOR and Portia consistently exhibit lower performance across all evaluation metrics and datasets compared to Pearson Correlation. PPCOR computes partial correlation matrices, which reflect conditional independence between variables, while Portia extends this approach by fitting linear regression models to infer directional regulatory relationships and break the inherent symmetry of precision matrices. Both methods rely on the assumption that the data follows a Gaussian distribution, as the sparsity patterns of precision and partial correlation matrices are valid indicators of conditional independence under this assumption. However, this assumption is likely inappropriate for single-cell RNA-seq data due to two key factors: (i) the high sparsity of the data, with many zero entries resulting from dropout events, could violate Gaussian properties; and (ii) gene expression distributions are often multimodal (in the statistical sense, where each cell type / cluster produces a mode), reflecting biological heterogeneity such as distinct cell states or types [26]. Simplifying these complex data distributions to a Gaussian framework may lead to improper modeling and biased inference, which could explain the observed performance decline in our metrics.

### Further evaluation of TF-gene directionality permutation

We excluded TF binding and Gene Set Recovery from the causal directionality analysis for a structural reason specific to GRN topology. In the original network, TFs are the source nodes, with edges pointing to a broad set of target genes (both TF and non-TF). When all edges are reversed, the only edges that still originate from a TF are those where the original target was itself a TF — i.e., only TF→TF edges survive as TF-sourced relationships. This fundamentally collapses the evaluation domain: TF binding and Gene Set Recovery are designed to assess how well a TF’s predicted target gene set matches binding evidence or pathway annotations, but after reversal, each TF’s outgoing neighborhood shrinks to only other TFs — a sparse, biologically distinct subset that is not representative of the full regulatory program. The metric is therefore no longer evaluating the same biological question; it has shifted from “does this TF regulate the right genes?” to “does this TF regulate the right other TFs?” — an inadvertent conceptual shift that would confound any directional sensitivity signal with a change in evaluation scope.

### Effect of skeleton filtering on GRN inference performance

To evaluate whether incorporating biological priors—such as known TF binding motifs—can improve GRN inference performance, we conducted an analysis in which we constructed dataset-specific skeletons based on motif and chromatin accessibility data, and used these priors to constrain inferred networks. We then assessed whether such prior-based filtering improves performance across our evaluation metrics.

To construct the skeletons, we combined TF motif information from ENCODE and JASPAR with two complementary sources of regulatory evidence for each dataset: (i) ATAC-seq peak–gene links, connecting open chromatin regions to target genes within ±150 kb, and (ii) promoter-proximal regions defined as ±1 kb around each gene’s transcription start site. This procedure was applied independently to each dataset, yielding skeletons comprising 13.5M edges (1,282 TFs, 17,075 genes) for OPSCA. Each inferred GRN was subsequently filtered to retain only skeleton-supported edges, and all evaluation metrics were recomputed on the pruned networks.

The results shows that the TF binding metric consistently increased after filtering for the skeleton (Supplementary Fig. 8-H), which is expected given that the skeleton is derived from the same motif databases underlying the TF binding metric, leading to a circularity effect. However, other metrics either do not change or show minor decrease following filtering. These findings support our main conclusion that incorporating prior biological information as a hard constraint can degrade GRN inference performance, likely due to the incompleteness and context-independence of current motif and accessibility annotations, which may exclude true context-specific regulatory interactions.

### CellOracle’s limitation in capturing enhancers

We observed notable peculiarities in CellOracle as an eGRN inference method. CellOracle uses Cicero to calculate peak co-accessibility, establishing a prior network that is subsequently used to prune the GRN inferred through regression models. While designed for eGRN inference, we have previously reported that CellOracle produces only promoter-based regulatory connections [98]. This limitation is further illustrated in Supplementary Fig. 8-G, where we annotate the types of CREs retained in the prior network obtained by CellOracle. Unlike other enhancer-based methods such as Scenic+, GRaNIE, and scGLUE—which yield a more balanced distribution of annotation types including *promoters, introns*, and *distal intergenic* regions—CellOracle retains only promoter-based connections. Given this restriction, our implementation of CellOracle in geneRNIB includes only promoters and skips the Cicero step to conserve computational resources. We encourage the authors to address this issue and revise the algorithm within geneRNIB to improve its handling of enhancer-based regulations.

For CRE annotation, we used the R package `ChIPseeker` [99] with parameters `tssregion` set to [− 1000, 1000] bp and `level` set to “`transcript`”. This step identifies the type of CREs and assigns them to different DNA regions, particularly `promoters`, `distal intergenic`, and `downstream (`≤ `300)`, with the latter two potentially acting as enhancers. `Distal intergenic` regions are located between genes, and `downstream` (≤ `300)` regions are found immediately after the end of a gene. Both types of regions can be involved in gene regulation.

## Supplementary Note 4

### Application-guided interpretation of our benchmark metrics

In our benchmark, we considered multiple metrics for holistic assessment of GRN inference performance. Nonetheless, each metric embodies distinct assumptions and captures complementary dimensions of GRN quality, which can guide model selection toward particular applications. The SEM and Virtual cell metrics are naturally aligned with non-linear perturbation modeling and can guide GRN model selection toward interpretable frameworks for drug response prediction and perturbation planning [47, 48]. In particular, the Virtual cell metric is well suited for personalized medicine applications, where large compound libraries can be screened computationally prior to experimental validation using only a patient’s baseline gene expression profile. The Regression metric, another predictive metric, differs from Virtual cell and SEM in that it evaluates whether inferred regulators can quantitatively predict their target genes’ expression levels (e.g., using linear models). This makes it particularly valuable for applications requiring identification of functional upstream controllers—for instance, determining which TFs regulate disease-relevant genes like IL-6 in autoimmune disorders [100]. Or, for cancer prognosis applications where a panel of oncogenic TFs (e.g. MYC [101]) measured from tumor biopsies can predict the expression of hundreds of downstream genes involved in apoptosis and proliferation, enabling risk stratification without whole-transcriptome sequencing.

The TF recovery metric, which evaluates whether a GRN can correctly identify which TF was perturbed based on downstream transcriptional effects, is particularly valuable in identifying the causal drivers of observed phenotypes. Models with high TF recovery scores can reliably pinpoint master regulatory TFs driving disease phenotypes—for instance, identifying key dysregulated factors in disease, or ranking candidate TFs for drug development where targeting upstream master regulators may be more effective than targeting individual downstream genes [102, 12].

The WS distance metric focuses on functional TF–gene relationships at individual edge resolution. This edge-level evaluation is especially informative for capturing regulatory interactions that are phenotypically important yet highly context-dependent. For example, in CAR-T cell engineering, the FOXO1→IL7R regulatory interaction is critical for T cell persistence: it is strong in naive T cells, where FOXO1 promotes IL7R-mediated survival signaling, but is weakened in exhausted T cells [103]. GRN models with high WS distance scores can accurately capture such context-dependent variations in regulatory strength, thereby informing about how to engineer CAR-T cells that preserve the FOXO1–IL7R axis and improve therapeutic durability.

Gene set recovery evaluates whether a model correctly identifies TFs that regulate pathways active in a given biological context, making it particularly valuable for applications that require pathway-level interpretability rather than precise edge-level reconstruction. For example, in drug repurposing screens where the goal is to determine how compounds modulate specific biological programs (e.g., JAK inhibitors suppressing inflammatory signaling [104]), models with strong Gene set recovery performance can accurately link relevant TFs to the affected pathways. This facilitates mechanistic interpretation of drug–pathway interactions by revealing the upstream regulatory factors driving the observed transcriptional response.

Together, these metrics help guide the selection of GRN inference methods by matching model strengths to the specific requirements of different biological applications.

### Perspectives on future developments of GRN inference methods

The findings of this study suggest that simpler algorithms—those with fewer underlying assumptions and preprocessing steps—often outperform more complex approaches. Multimodal GRN inference methods that integrate TF motif databases can inadvertently introduce biases if motif information is not properly contextualized to the biological setting under study. These limitations could arise both from the integration methodology itself and from intrinsic constraints of the underlying databases.

Regarding motif resources, although it is often stated that a large fraction of TFs is covered, this coverage is neither exhaustive across TFs nor comprehensive across their binding motifs [105, 12]. Moreover, motifs are derived from a mixture of experimental assays and computational inference rather than being uniformly obtained from direct, TF-specific binding measurements [105, 12]. As a result, they may reflect incomplete or partially transferred binding preferences. Even when a TF is represented in a motif database, the annotated motifs often fail to capture the full diversity of its binding behavior, including context-dependent binding, cofactor-mediated interactions [106, 107, 108], chromatin-dependent accessibility effects [109, 110], and alternative or low-affinity binding configurations [111, 112]. In addition, these databases are largely context-agnostic, having been compiled across diverse cell types, tissues, and experimental conditions, which can further obscure condition-specific regulatory logic [105, 12]. Taken together, while these resources remain valuable for guiding inference and reducing search space, their effective use requires careful consideration of biological context and integration strategy to avoid over-reliance on potentially non-representative motif priors.

A more effective approach may be to reverse this workflow: first construct a preliminary GRN from prior knowledge, then refine it further using expression data to contextualize interactions. For example, an initial network could encode global regulatory interactions from motif datasets, subsequently fine-tuned with expression and chromatin accessibility data to capture context-specific regulation. Graph neural networks (GNNs) are well-suited for this task, leveraging prior knowledge to establish initial connectivity while optimizing relationships in latent space [113].

Similarly, foundation models—such as those leveraging large language models (LLMs)—use vast datasets to learn global regulatory patterns, which can then be contextualized for specific datasets. However, our evaluation of three foundations methods—scGPT, `scFoundation`, and scPRINT—suggests that these models struggle to effectively contextualize GRNs, often underperforming compared to methods based solely on expression data. Despite this limitation, foundation models hold significant promise. Refining their contextualization capabilities (potentially through transfer learning) could significantly enhance their utility. geneRNIB provides an ideal framework for benchmarking these advances, offering systematic evaluation across diverse datasets. Notably, there is currently no foundation model that explicitly contextualizes GRNs using multiomics data, presenting a compelling direction for future research.

Interventional data offer another promising avenue for improving GRN inference. CausalBench highlighted substantial gains when incorporating such data [8]. For example, `BetterBoost`, an extension of `GRNBoost` integrating perturbation data and causal structure, ranked among the top-performing methods in their benchmark. While these approaches are effective for GRN inference when interventional data is available, their broader applicability is limited to observational data. Expanding evaluations using geneRNIB, which offers a broader range of datasets and metrics than CausalBench, could help clarify their utility.

Finally, while most GRN inference methods focus on transcriptional regulation inferred from expression and chromatin accessibility data, gene regulatory mechanisms go beyond these two modalities. As multiomics integration advances, GRN inference strategies must evolve accordingly, incorporating additional layers of regulation to fully exploit the richness of high-dimensional biological data.

### Guidance on future integration of inference and evaluation datasets

In the absence of actual ground truth, we believe that large-scale perturbation datasets are key to establishing a robust benchmark for GRN inference. For this approach to be effective, the design of inference and evaluation datasets should consider several critical aspects.

Most GRN inference methods rely on gene co-expression or predictive patterns to propose or refine putative regulatory interactions. Therefore, inference datasets must exhibit sufficient transcriptional variability to enable reliable network reconstruction. In the majority of our datasets, this variability is primarily introduced through experimental perturbations. To avoid information leakage and overly optimistic performance estimates, we carefully split perturbation datasets into distinct inference and evaluation subsets (Methods), ensuring that identical perturbation conditions are not shared between the two. In some settings, however, GRN inference can be performed on a complementary dataset, provided that it is contextually aligned with the experimental conditions used for evaluation. For example, the OPSCA dataset represents a scenario in which the dominant source of transcriptional variability arises from differences in cell type rather than from perturbations. In such cases, cell-type heterogeneity can serve as an alternative driver of regulatory signal for network inference. An alternative strategy involves inferring GRNs directly from the perturbation dataset and evaluating them on held-out cell types, as demonstrated by Kamal et al. [5].

However, a general limitation of current approaches is that individual cell types and perturbations—particularly broad-acting interventions such as drug treatments—may induce context-specific regulatory programs. While such experiments often share higher-level regulatory mechanisms and are expected to preserve core network structure, nuanced regulatory interactions may be diluted or missed when GRNs are assessed under aggregated conditions. To mitigate this limitation, we diversified our evaluation by including multiple types of perturbations, such as genetic knockout datasets with more localized regulatory effects, as well as temporal datasets in which GRN inference was performed at one time point and evaluation at another. By assessing GRN performance across these diverse scenarios and observing consistent trends, we believe that the impact of this limitation on our conclusions is likely limited.

For multiomics datasets, additional complexities arise, as they integrate multiple layers of information, such as transcriptomics, epigenomics, and proteomics, each capturing distinct but complementary regulatory features. Perturbations can impact not only gene expression but also these additional layers. Therefore, a fully context-specific evaluation must consider all these layers simultaneously, with measurements captured under identical experimental conditions. In our integration so far, we assumed that the baseline multiomics data is fully relevant to the perturbational settings. However, this assumption has limitations, as perturbations could also alter chromatin accessibility or other epigenetic features. Unfortunately, we could not identify a dataset that comprehensively addresses these requirements. To overcome this limitation, geneRNIB supports dynamic updates of datasets, allowing future integration of such datasets once they are produced.

Currently, our integration focuses on multiomics datasets combining scRNA-seq and scATAC-seq data. While valuable, these alone are insufficient to fully characterize the complexity of gene regulation. Coupled multiomics profiling approaches with more modalities are particularly promising in addressing this complexity [2]. For instance, NEAT-seq profiles intra-nuclear proteins, chromatin accessibility, and gene expression; scChaRM-seq captures DNA methylation, chromatin accessibility, and gene expression; ATAC-STARR-seq links chromatin accessibility to functional enhancer activity; and Phospho-seq measures protein phosphorylation alongside chromatin and transcriptional states. Each additional modality provides complementary evidence that would otherwise be simplified in GRN reconstruction. For example, many GRN inference methods associate TFs with target genes based solely on gene expression [12, 3, 5, 19], overlooking the nonlinearity between mRNA and protein levels. Incorporating protein level data can address this limitation by providing direct measurements of TF protein levels, thereby enhancing the accuracy and biological relevance of GRN inference. Similarly, while chromatin accessibility measured by peak data is typically associated with functional regions, epigenetic properties such as DNA methylation, histone modifications (e.g., acetylation, methylation, phosphorylation), and chromatin remodeling factors can further refine and modulate the role of chromatin state in gene regulatory mechanisms [114, 2].

Time-series measurements provide an additional layer of valuable information, holding the potential to substantially guide the inference process. In our study, the 300BCG and SoundLife datasets contain temporal data, where we used the perturbation samples from earlier measurements for GRN inference and subsequent measurements for evaluation, assessing whether the regulatory relationships obtained earlier are applicable to the more stable system measured later. The results demonstrated that this is indeed the case (Supplementary Fig. 1), where the evaluation showed a similar trend of comparative performance of GRN inference methods to other static datasets. This indicates that the inferred regulatory relationships hold true across different temporal states.

However, it is important to note that our analysis so far has only considered different time points for inference and validation separately, treating each time point as static data, rather than leveraging the temporal dynamics for GRN inference. In our approach, the system is assumed to have reached equilibrium at the time of measurement. True temporal information, by contrast, introduces dynamic aspects that offer insights into gene and transcription factor interactions over time, potentially enabling more robust causal inference (e.g., Granger causality). Multiple GRN inference methods have been developed specifically to leverage such temporal data, either in the form of time series or pseudotime trajectories [115, 116]. However, pseudotime trajectories, reconstructed *post hoc* without direct time measurements, do not fully capture the true temporal dynamics of a system. Meanwhile, time-resolved measurements, while offering higher fidelity, are costly and logistically challenging, particularly in large-scale perturbation studies. An alternative strategy is RNA metabolic labeling, which provides an intermediate level of time resolution by distinguishing newly synthesized transcripts from pre-existing ones [117]. This approach enables inference of regulatory activity over shorter timescales without requiring full longitudinal sampling. As time-series datasets become more accessible, extending geneRNIB to integrate and leverage these datasets represents a promising avenue for advancing GRN inference methods and enabling broader, more context-specific applications.

## Footnotes

1 https://openproblems.bio/

2 https://scglue.readthedocs.io/

3 https://scenicplus.readthedocs.io/en/latest/pbmc_multiome_tutorial.html

4 https://buenrostrolab.github.io/FigR/articles/FigR_shareseq.html

5 https://morris-lab.github.io/CellOracle.documentation/

6 https://grp-zaugg.embl-community.io/GRaNIE/articles/GRaNIE_singleCell_eGRNs.html

7 https://arboreto.readthedocs.io/en/latest/

8 https://pyscenic.readthedocs.io/en/latest/tutorial.html

9 https://www.rdocumentation.org/packages/PPCOR/versions/1.1/topics/pcor

10 https://github.com/AntoinePassemiers/Portia

11 https://github.com/cantinilab/scPRINT/

